# A synaptic wiring diagram underlying modality-specific ascending pathways of the superficial spinal dorsal horn

**DOI:** 10.64898/2026.09.27.754843

**Authors:** Wangchu Xiang, Ramin Khajeh, Kevin Delgado, Jingyi Liu, Adrian Holtrup, Qiyu Zhang, Sidrick Cameron, Tri Nguyen, J. Alex Bae, Akhilesh Halageri, Nico Kemnitz, Thomas Macrina, Borja Moya Llamas, Alice Schildkamp, Mohd Freezely Mazri, Julianne Ni, Ruairi Roberts, Laia Serratosa Capdevila, Ruoming Wang, Amelia Dsouza, Aliya Norton, Songyang Wang, Dong Zhang, Paul Lindu Tang, Seungwon Choi, Wei-Chung Allen Lee, David D. Ginty

**Affiliations:** Department of Neurobiology, Harvard Medical School, 220 Longwood Avenue, Boston, MA, USA; Howard Hughes Medical Institute, Harvard Medical School, 220 Longwood Avenue, Boston, MA, USA; Zetta AI, Sherrill, NJ, USA.; Aelysia LTD, Bristol, UK; Department of Psychiatry, University of Texas Southwestern Medical Center, Dallas, TX, USA; FM Kirby Neurobiology Center, Boston Children’s Hospital, Boston, MA

## Abstract

The spinal cord dorsal horn transforms tactile, thermal, pruritic, and nociceptive signals from peripheral somatosensory neurons into outputs to the brain, but how dorsal horn circuits organize diverse inputs into modality-selective ascending pathways is poorly understood. Using dSC1, a large-scale electron microscopy dataset of the mouse dorsal horn, we mapped the circuitry connecting primary sensory afferents to projection neurons of the anterolateral tract (ALT), a principal pathway conveying somatosensory information to the brain. We find that afferents encoding distinct modalities are routed onto connectivity-defined populations of ALT projection neurons. Nociceptors and cold thermoreceptors directly innervate distinct lamina I ALT projection neuron populations specialized for conveying noxious and cooling signals, respectively, whereas tactile and pruritic signals instead pass through a recurrent, multilayered network involving interneurons called vertical cells before reaching a separate ALT projection population. This distinction, direct versus polysynaptic wiring, underlies marked differences in sensory convergence and establishes segregated ascending channels for conveying signals encoding the principal somatosensory modalities. Together, our findings define an architectural logic by which the dorsal horn integrates and routes tactile, thermal, pruritic, and nociceptive information into distinct ALT output pathways to the brain.

## Main

Mammals encounter a range of somatosensory stimuli, including touch and its submodalities (pressure, vibration, skin stretch, and hair deflection), cold, heat, and chemicals. These stimuli govern reflexes, including protective nocifensive responses and scratching, and contribute to cognition and behaviors, including social interactions and communication. Somatosensation is enabled by more than a dozen physiologically distinct dorsal root ganglion (DRG) sensory neuron subtypes that innervate the skin and other body organs and respond to select mechanical, thermal, and chemical stimuli (Abraira and Ginty, 2013; Julius, 2013; Lay and Dong, 2020; Lumpkin and Caterina, 2007). These low-threshold mechanoreceptors (LTMRs), high-threshold mechanoreceptors (HTMRs), thermoreceptors, polymodal sensory neurons, and nociceptors form synapses with interneurons and, in some cases, ascending projection neurons in the spinal cord dorsal horn (Abraira and Ginty, 2013; Abraira et al., 2017; Braz et al., 2014; Chen et al., 2024; Choi et al., 2020; Morin, 1955; Todd, 2010; Wang et al., 2022), thus rendering the dorsal horn the earliest stage of somatosensory information processing.

Laminae I-II of the dorsal horn receives inputs from a range of DRG sensory neuron subtypes and serves as a major sensory processing center for tactile, thermal, and chemical cues (Braz et al., 2014; Koch et al., 2018; Qi et al., 2024; Ran et al., 2016; Ross, 2011). While lamina I contains interneurons as well as projection neurons that relay sensory information to the brain via the anterolateral tract (ALT), lamina II (substantia gelatinosa) contains only interneurons and is devoid of ascending projection neuron somata (Chen et al., 2024; Han et al., 1998; Lahuerta et al., 1990)(**Figure 1a**). The number and diversity of somatosensory neuron (Bhuiyan et al., 2024; Qi et al., 2024; Usoskin et al., 2015), dorsal horn interneuron, and projection neuron subtypes (Gatto et al., 2019; Roome et al., 2026; Russ et al., 2021), together with their dense and intermingled processes and synapses, have made it challenging to define modality-specific sensory processing circuits of laminae I-II. To overcome these challenges, here we used a large-scale EM volume of the mouse lumbar spinal cord dorsal horn (the dSC1 dataset) and a SPinal cord Integrated NEuron Classifier (SPINE) (Xiang et al., 2026a) to map synaptic connections between physiologically distinct sensory neuron subtypes and neurons of dorsal horn laminae I-II. We focused on laminae I-II excitatory interneurons (ExNs) and projection neurons (PNs) to begin to understand how the superficial dorsal horn circuitry transforms somatosensory neuron signals into ascending output streams to the brain via lamina I ALT projection neurons.

**Figure 1.**
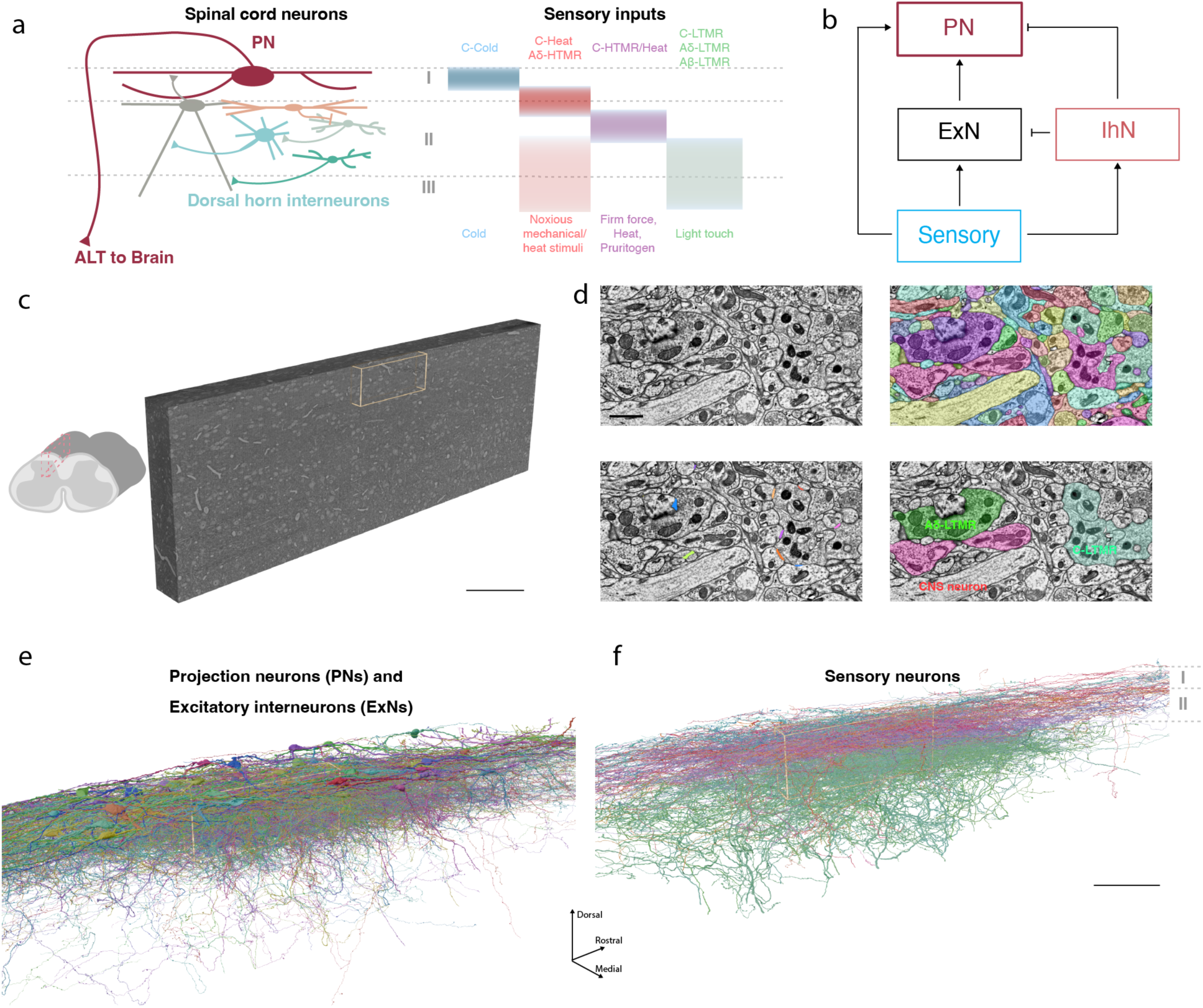
Serial EM reconstruction of sensory processing circuits of the spinal cord dorsal horn. **a**, Schematic of dorsal horn sensory circuits. Local interneurons and projection neurons (left) integrate input from distinct primary afferent subtypes that convey different cutaneous sensory modalities and terminate in specific dorsal horn laminae (right). Lamina I projection neurons send ascending axons via the anterolateral tract (ALT) to the brain. **b**, Simplified summary of circuit organization: primary afferent input and local excitatory (ExN) and inhibitory (IhN) interneurons shape PN output. **c**, Parasagittal section of the lumbar spinal cord showing approximate location of the acquired EM volume. The yellow box marks the region in which all ExNs were reconstructed. Additional PNs and Vertical cells outside the yellow box were also reconstructed (see text). Scale bar, 200 µm. **d**, Automated analysis pipeline. Neuronal processes were segmented using CNN-based methods (top right, segments shown in distinct colors), synapses were automatically predicted across the EM volume (bottom left, clefts highlighted in distinct colors), and sensory afferent subtypes were classified using SPINE (bottom right, subtypes overlaid with distinct colors). Scale bar, 1µm. **e**, 3D rendering of reconstructed 177 dorsal horn excitatory interneurons and 46 projection neurons. Different cells are colored randomly. **f**, 3D renderings of 159 reconstructed primary sensory afferents: C-Cold neurons (blue), CGRP^+^ C-nociceptors and Aδ-HTMRs (orange), C-HTMR/Heat neurons (purple), and C-LTMRs, Aδ-LTMRs, and Aβ-LTMRs (green). Scale bar, 100 µm (e,f).

### Reconstruction of excitatory neurons in laminae I-II

To map sensory neuron-to-ALT output pathways of the superficial dorsal horn, we examined synaptic connectivity between sensory neuron subtypes, excitatory and inhibitory lamina I-II interneurons (ExN and IhN), and lamina I ALT projection neurons (PNs) within the dSC1 EM volume (**Figure 1b,c**). This EM volume spans 1.5 mm along the rostral–caudal axis, 190 µm along the medial–lateral axis, and 600 µm along the dorsal–ventral axis of the lumbar L3-L4 mouse spinal cord dorsal horn, allowing for reconstruction of complete central axonal arbors of primary afferents as well as the dendritic arbors and axons of dorsal horn neurons. Neurons in the EM volume were automatically segmented, and synapses were identified using convolutional neural networks (CNNs). To identify the synapses of physiologically defined sensory neuron subtypes onto spinal cord neurons, we developed the SPINE classifier that accurately predicts DRG neuron subtypes directly from EM images (Xiang et al., 2026a). This approach enabled sensory neuron subtype-resolved reconstruction of neuronal circuits of the dorsal horn (**Figure 1d**).

We reconstructed the axons and dendrites of all ExNs that have their somata within a central region of interest (230 × 110 × 31 µm along the rostral–caudal, dorsal–ventral, and medial– lateral axes, respectively; yellow bounding box in **Figure 1c,e,f**), yielding 100 fully reconstructed ExNs spanning laminae I-II (**Figure S1c**). To extend our sampling beyond the central ROI and capture additional ExNs that may participate in feedforward pathways to lamina I ALT PNs, we additionally reconstructed a population of vertical cells (**Figure S1a,b,d**), which are ExNs proposed to convey signals from lamina II/substantia gelatinosa to lamina I ALT PNs (Braz et al., 2014; Duan et al., 2014; Lu and Perl, 2005). We also reconstructed 46 lamina I ALT PNs distributed over 1.2 mm along the rostral–caudal axis (**Figure S1e**). In total, our dataset comprises 177 excitatory interneurons (100 ExNs and 77 vertical cells) and 46 lamina I ALT PNs (**Figure 1e; Figure S1f**).

The 177 ExNs had axons that predominantly extended along the rostral–caudal axis within laminae I-II. Notably, 74% of the ExNs additionally had a single ventrally directed axon that descended beyond lamina IV before exiting the volume, most often in the medial–lateral direction (**Figure S1c**), indicating that many superficial dorsal horn ExNs form synapses upon neurons beyond the superficial laminae and along the medial–lateral axis. In total, the 177 reconstructed ExNs formed 38,938 synapses within the EM volume, of which 84.5% were located within laminae I-II (**Figure S1h**). Because many ventrally directed axons exited the EM volume, we could not quantify the full complement of synapses made by these projections (but see Discussion). This densely reconstructed population of ExNs and PNs provides a platform for defining feedforward excitatory circuit motifs linking distinct classes of primary afferents to dorsal horn interneurons and lamina I ALT PNs. In a companion study (Xiang et al., 2026a), we reconstructed the axonal arbors of 161 primary sensory afferents (**Figure 1e**) together with a subset of inhibitory interneurons (IhNs) that mediate sensory-evoked presynaptic inhibition. Combined, these reconstructed datasets allowed us to visualize and quantify synapses between sensory neuron subtypes, ExNs, IhNs, and PNs.

### Defining interneuron subtypes by sensory synaptic inputs and morphology

Prior work characterizing laminae I-II interneurons described several morphologically defined interneuron subtypes (Grudt and Perl, 2002; Heinke et al., 2004; Todd and Lewis, 1986; Yasaka et al., 2007). Indeed, classic neuroanatomical studies, including those of Cajal, Perl and colleagues, identified spinal cord interneuron subtypes such as vertical and radial cells (Braz et al., 2014; Gobel, 1978; Grudt and Perl, 2002; Lu and Perl, 2005; Maxwell et al., 2007; Ramón y Cajal, 1909) that can be readily recognized by their dendritic morphologies. However, many interneurons in laminae I-II do not readily fall into the canonical morphological categories (Grudt and Perl, 2002). We therefore sought to classify reconstructed laminae I-II excitatory interneuron (ExN) subtypes based on their dendritic morphology and synaptic inputs from primary somatosensory neurons. For this, we applied the SPINE model (Xiang et al., 2026a) to identify and quantify the synapses from different sensory neuron subtypes on the 177 reconstructed laminae I-II ExNs (**Figure 2a,b; Figure S2a**). We then integrated these synaptic input features with morphological metrics extracted from neuron skeletons to embed and cluster the ExNs (**Figure S2b; Methods**). This analysis yielded six laminae I-II ExN clusters that differ in dendritic morphology and synaptic input composition from the different sensory neuron subtypes (**Figure 2d**).

**Figure 2.**
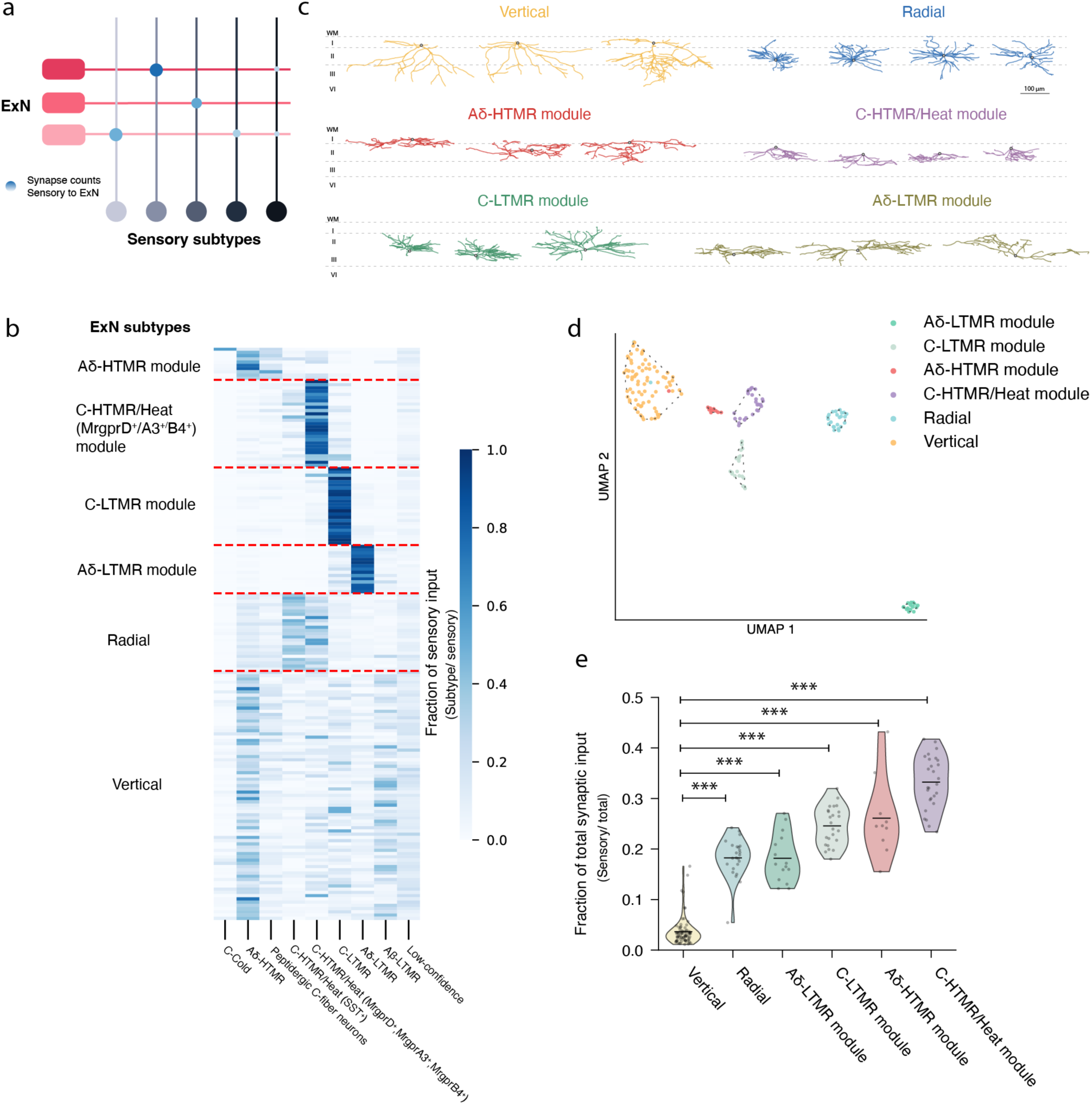
Sensory input to laminae I-II excitatory interneurons. **a**, Schematic of the sensory neuron–to–ExN connectivity matrix. Sensory neuron subtypes (columns) are color-coded, and ExN types (rows) show distinct input patterns. **b**, Fraction of sensory input from each afferent subtype (columns) onto each ExN group (rows). **c**, Representative skeleton reconstructions of ExNs for each subtype. **d**, UMAP embedding of ExNs using both synaptic input features and dendritic morphology features. Dashed outlines indicate HDBSCAN-identified clusters (Methods); colors denote the final subtype assignments. **e**, Violin plots showing the fraction of sensory versus non-sensory synaptic inputs across the six ExN clusters. (Vertical versus radial, p = 2.97×10^-6^; vertical versus Aδ-LTMR module, p = 2.14×10^-4^; vertical versus C-LTMR module, p = 2.74×10^-13^; vertical versus Aδ-HTMR module, p = 7.06×10^-7^; vertical versus C-HTMR/Heat module, p = 1.81×10^-24^, using a Kruskal–Wallis test, and Dunn posthoc test with Holm–Bonferroni-correction. Vertical cells, n=77; radial cells, n=24; Aδ-LTMR module ExNs, n=15; C-LTMR module ExNs, n=24; Aδ-HTMR module ExNs, n=10; C-HTMR/Heat module ExNs, n=27.)

Two lamina I-II ExN clusters corresponded to canonical morphologically defined interneuron subtypes with tight laminar localization: vertical cells, with somata near the lamina I-II border, and radial cells, with somata in lamina IIid (**Figure 2c**). Strikingly, vertical cells received far fewer sensory neuron synapses than the other five cell type clusters. Fewer than 5% of synaptic inputs to vertical cells arose from sensory neurons, whereas all other ExN cell clusters received significantly higher proportions of monosynaptic sensory neuron input (**Figure 2e**). Vertical cells derive most of their limited sensory inputs from Aδ-HTMRs, with a subset also receiving monosynaptic Aβ-LTMR input. In contrast, a second cluster, comprised of radial cells, received on average ∼18% of their synaptic inputs from sensory neurons, preferentially C-HTMR/Heat (Sst^+^) neurons, while also receiving inputs from other C-HTMR/Heat subtypes (MrgprD^+^, MrgprB4^+^, MrgprA3^+^).

The other four ExN clusters had a large fraction (18%-33%) of their synaptic inputs from sensory neurons, with each one preferentially receiving monosynaptic input from a single major afferent class. We refer to these four ExN clusters as sensory modules, named by the sensory neuron subtype they preferentially sample. ExNs in the Aδ-HTMR module received 43.7% ± 19.7% of their sensory input from Aδ-HTMRs; those in the C-HTMR/Heat module received 69.5% ± 18.0% from C-HTMR/Heat afferents (MrgprD^+^, MrgprB4^+^, MrgprA3^+^); those in the C-LTMR module received 82.6% ± 10.0% from C-LTMRs; and those in the Aδ-LTMR module received 77.1% ± 11.4% from Aδ-LTMRs (**Figure 2b; Figure S2c**). While neurons in these ExN modules exhibited diverse morphologies, many could be broadly categorized by classical criteria as central cells (Grudt and Perl, 2002; Maxwell et al., 2007) (**Figure 2c**). The somata locations of neurons within these four ExN modules showed a dorsoventral trend that broadly mirrored the termination patterns of the sensory afferents they preferentially sample, while exhibiting substantial spatial overlap between the modules (**Figure S2d**). Notably, even within the same laminar region, neurons exhibited strong module-specific input preferences, indicating that sensory input identity cannot be explained by soma position alone.

In summary, lamina I-II ExN neurons can be classified based on a combination of sensory neuron synaptic input profiles and dendritic morphology. Most ExN subtypes receive strong monosynaptic inputs from a single afferent class, suggesting an early, modular organization that preserves distinct sensory channels. In contrast, vertical cells receive predominantly CNS-derived inputs, consistent with their proposed role as third- or higher-order neurons in feedforward pathways to PNs (Braz et al., 2014; Duan et al., 2014; Lu and Perl, 2005). We observed little Aβ-LTMR input across the reconstructed ExNs, with only weak input detected in a subset of vertical cells. This likely reflects the focus of our reconstruction on neurons with somata in laminae I-II, whereas Aβ-LTMR afferents terminate predominantly in more ventral regions of the dorsal horn. Accordingly, we did not further investigate Aβ-LTMR processing pathways in this study. Future studies analyzing deeper dorsal horn laminae will resolve the spinal circuits underlying Aβ-LTMR processing.

### Defining lamina I ALT projection neuron subtypes by their sensory inputs

Classic electrophysiological recording studies reported substantial diversity in lamina I ALT PN tuning, including PNs that selectively respond to noxious stimuli or cold temperature as well as polymodal PNs that respond across stimulus modalities and intensities (Christensen and Perl, 1970; Ferrington et al., 1987; Han et al., 1998; Light et al., 1993; Price et al., 1978). More recently, genetic access to molecularly defined ALT PN subtypes (e.g., Tacr1^+^ and Gpr83^+^) has suggested that morphologically and functionally distinct ALT PNs relay modality-specific signals to the brain (Ahanonu et al., 2024; Choi et al., 2020; Razlan et al., 2025; Warwick et al., 2022; Yarmolinsky et al., 2025; Zhang et al., 2026). Therefore, as with the laminae I-II ExNs, we used both morphological and SPINE-assisted sensory neuron synaptic analyses to classify lamina I ALT PNs.

Previous work showed a bimodal distribution of soma size among lamina I neurons, with PNs having larger somata than interneurons (Al Ghamdi et al., 2009; Cameron et al., 2015; Cheunsuang and Morris, 2000). We examined soma size of 456 lamina I neurons in our EM dataset and confirmed this bimodal distribution (**Figure S3a**). Using soma size as a criterion, we identified 46 putative lamina I PNs, representing approximately 10% of the lamina I neurons examined, consistent with previous estimates that PNs comprise 5–10% of neurons in this region (Al Ghamdi et al., 2009; Bice and Beal, 1997a, b). Putative PNs had significantly larger somata than lamina I interneurons (1800.9 ± 365.5 μm³ vs. 787.3 ± 269.0 μm³) and exhibited more elaborate dendritic arbors (**Figure S3b,c**), occupying on average a threefold larger dendritic territory than interneurons (**Figure S3d**). This expanded dendritic morphology is consistent with previous descriptions of lamina I PNs (Lima and Coimbra, 1986; Spike et al., 2003). Among these, six PNs exhibited a distinctive morphology characterized by one or more ventrally directed dendrite that extended into the LTMR-recipient zone, with some dendrites reaching as deep as lamina V (**Figure 3a**). This morphology is consistent with reported features of Gpr83^+^ lamina I PNs (Choi et al., 2020). To further test this correspondence, we sparsely genetically labeled Gpr83^+^ ALT PNs and confirmed that a subset of labeled Gpr83^+^ PNs have prominent, ventrally projecting dendrites that extend as deep as lamina V (**Figure S4**). We therefore grouped these six PNs with ventrally projecting dendrites as P1 (putative Gpr83^+^ PNs). The remaining 40 PNs were referred to as P2; these neurons exhibited dendritic arbors confined to lamina I and IIo, consistent with the morphology of sparsely labeled lamina I Tacr1^+^ PNs (**Figure S4b)** (Choi et al., 2020).

**Figure 3.**
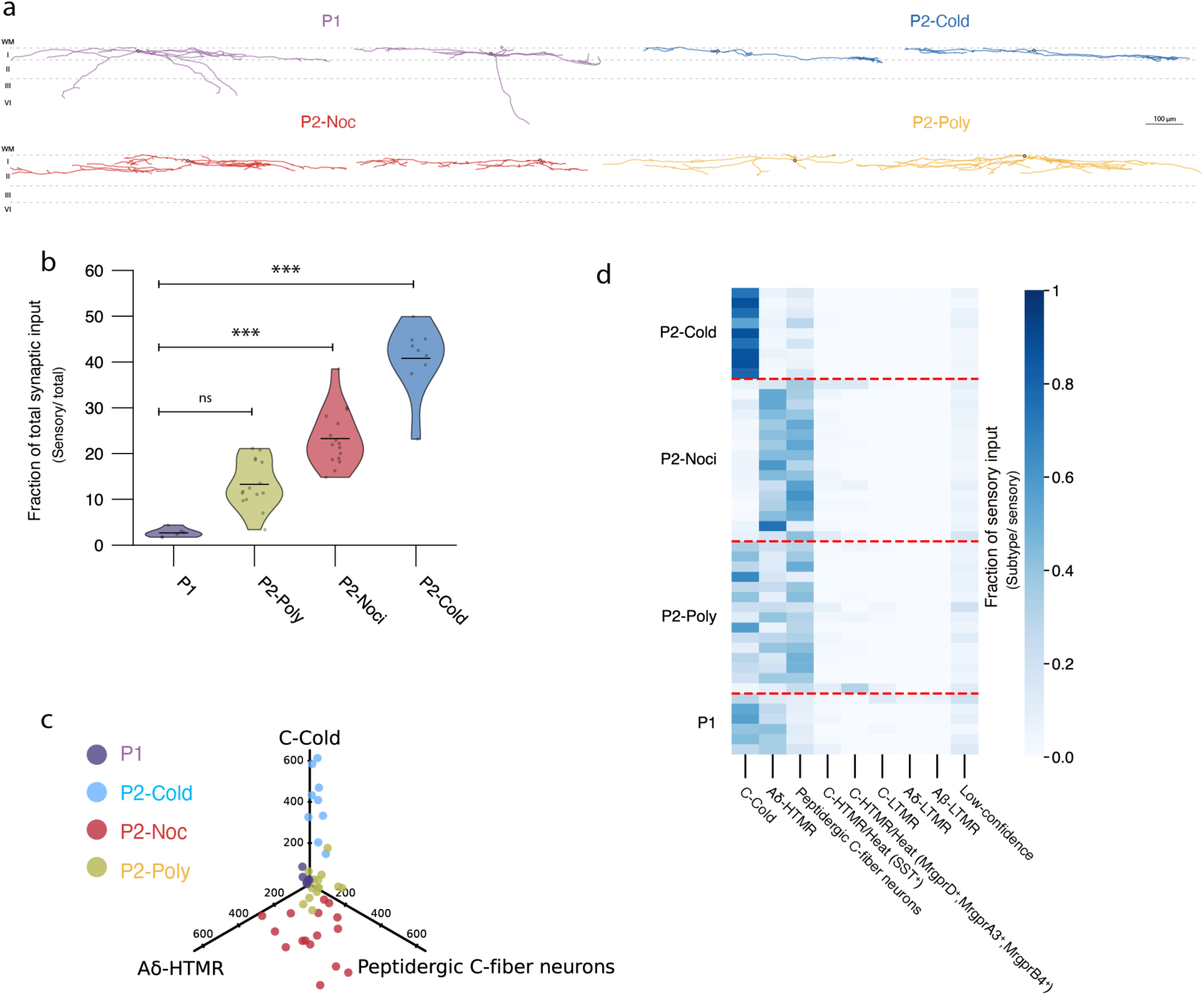
A wide variation of monosynaptic sensory neuron input to laminae I anterolateral tract projection neurons. **a**, Representative skeleton reconstructions of the four connectivity-defined PN subtypes. **b**, Violin plots showing the fraction of sensory neuron versus non-sensory neuron synaptic inputs across the four PN clusters. (P1 versus P2-Poly, p = 0.072; P1 versus P2-Nociceptor, p = 4.11×10^-4^; P1 versus P2-Cold, p = 6.62×10^-7^, using a Kruskal–Wallis test, and Dunn posthoc test with Holm–Bonferroni-correction. P1 neurons, n=6; P2-Cold neurons, n=9; P2-Nociceptive neurons, n=16; P2-Polymodal neurons, n=15.) **c**, PN subtypes stratified by sensory drive from three major sensory neuron groups that monosynaptically connect to PNs (C-Cold neurons, Aδ-HTMRs, and peptidergic C-fiber neurons). Each dot represents one PN. Each axis represents the absolute number of synapses from one of the three sensory neuron subtypes. **d**, Fraction of synaptic inputs derived from each sensory neuron subtype (columns) onto each PN group (rows). Note that the number of sensory neuron synapses onto P1 PNs is very low compared to the P2 PN subtypes (panel c).

We next used SPINE to quantify sensory afferent synaptic inputs onto the 46 PNs to further classify these neurons. Strikingly, P1 neurons received few direct monosynaptic sensory inputs (2.7 ± 1.0% of total synapses), whereas P2 neurons received substantially more direct sensory input (**Figure 3b**). We therefore embedded P2 PNs in sensory input feature space and clustered them based on their afferent input composition (**Figure S3g**). This analysis subdivided P2 neurons into three groups with distinct patterns of C-Cold neuron and nociceptor input (**Figure 3c,d; Figure S3f**), which we refer to as P2-Cold, P2-Nociceptive, and P2-Polymodal PNs. P2-Cold PNs received the highest proportion of monosynaptic sensory input, with 40.8 ± 7.5% (mean ± SD) of their total synaptic input arising directly from sensory neurons and C-Cold afferents accounting for 81.7 ± 6.1% of their sensory inputs. P2-Nociceptive PNs received 23.3 ± 6.1% of their total synaptic input directly from sensory neurons and preferentially sampled nociceptor inputs from Aδ-HTMRs and peptidergic (CGRP^+^) C-fiber neurons, which together comprised 87.1 ± 5.1% of their sensory inputs. In contrast, P2-Polymodal PNs received a smaller proportion of their total synaptic input directly from sensory neurons (13.3 ± 5.2%) but integrated inputs from sensory neuron subtypes encoding multiple modalities, including C-Cold neurons (27.6 ± 18.9% of sensory inputs), peptidergic C-fiber neurons (36.1 ± 11.7%), and Aδ-HTMRs (19.1 ± 9.8%).

These findings suggest that lamina I PNs are organized into four principal output channels and provide a circuit basis for the long-standing observation that lamina I PNs exhibit diverse tuning properties—including cold-selective, nociceptive-selective, and polymodal ALT PN subtypes (Christensen and Perl, 1970; Han et al., 1998). These results are also consistent with recent reports of molecularly defined PN subtypes such as Gpr83^+^ and Tacr1^+^ PN populations (Choi et al., 2020) and a Calb1^+^ cold-selective ALT PN (Razlan et al., 2025).

### Two connectivity defined vertical cell subtypes

ALT PNs represent the ascending output of superficial dorsal horn circuits and most of their synaptic inputs are from CNS neurons (i.e. non-sensory), which suggests that their tuning properties are likely shaped by inputs from local interneurons. Therefore, we next analyzed synaptic connectivity between the 177 reconstructed lamina I-II ExNs (ExN-ExN synapses) and between the 177 ExNs and the 46 lamina I PNs (ExN-PN synapses).

Beginning with vertical cells, our analysis revealed an unexpected finding that vertical cells readily segregate into two populations, which we term vertical 1 (Vert1) and vertical 2 (Vert2) neurons (**Figure S5a,b**). Vert1 neurons received strong excitatory input from radial cells and Vert 2 neurons, and they also formed robust recurrent connections with other Vert1 neurons. In contrast, Vert2 neurons received substantial excitatory input from all four sensory modules but showed minimal input from radial cells and other vertical cells. Despite their distinct synaptic inputs from other ExNs, Vert1 and Vert2 neurons exhibited similar ventrally projecting dendrites and overall sensory neuron input profiles (**Figure S5c**) although, as noted above, monosynaptic sensory input was very sparse for both groups (2.7% and 4.7% respectively), with Vert2 neurons exhibiting modestly higher direct sensory inputs, including more Aδ-HTMR synapses (**Figure S5d,e**).

This connectivity-based division of vertical cells into two subtypes aligns with recent reports of two molecularly defined vertical cell populations, which are marked by NPFF and GRPR and differ in morphological and electrophysiological properties (Mu et al., 2017; Polgar et al., 2023; Quillet et al., 2023). Based on these parallels, we hypothesized that the two connectivity-defined vertical cell subtypes correspond to NPFF^+^ and GRPR^+^ vertical cells. A prior study reported that NPFF^+^ vertical cells exhibit higher dendritic spine density than GRPR^+^ vertical cells (Quillet et al., 2023). We confirmed this finding using sparse genetic labeling of NPFF^+^ and GRPR^+^ vertical cells and found that Vert2 and NPFF^+^ neurons have more spines compared to Vert1 and GRPR^+^ neurons (**Figure S6**). We therefore designated Vert2 neurons as putative NPFF^+^ vertical cells and Vert1 neurons as putative GRPR^+^ vertical cells. Consistent with this, GRPR^+^ vertical cells have been reported to form recurrent connections (Polgar et al., 2023), matching our observation that Vert1 neurons uniquely exhibit an abundance of recurrent synaptic connections. Together, these findings establish two vertical cell populations defined by their connectivity, Vert1 and Vert2 neurons, which correspond to genetically defined vertical cell subtypes.

### A lamina I/II neuron wiring diagram

We next asked how sensory neuron signals are routed to PNs, either through direct monosynaptic connections to PNs or through indirect, polysynaptic connections to PNs involving ExNs. Thus, we generated synaptic connectivity matrices for the sensory neuron subtypes and the synaptically defined ExNs and PNs (**Figure 4a,b**). In addition, to assess the extent to which spatial proximity between presynaptic axons and postsynaptic dendrites could account for synapse number, we also quantified the corresponding axon–dendrite overlap for each pairing (**Figure 4a,c**). This measurement allowed us to identify neuron subtype pairs that were preferentially connected relative to their opportunity for forming synaptic contacts (**Figure 4a,c; Figure S7f**) (Ding et al., 2025; Kuan et al., 2024; Lee et al., 2016). The resulting synapse-density matrix between lamina I/II ExNs and lamina I PN subtypes was highly structured and could not be explained by axon– dendrite overlap or spatial location alone. For example, radial cell axons and Vert1 dendrites showed only modest overlap but formed many synapses, resulting in a high density of synapses between these two cell types (**Figure 4b,c; Figure S7a**). Conversely, Aδ-LTMR module neuron axons exhibited a high degree of overlap with the dendrites of other Aδ-LTMR module neurons, yet few synapses were observed between Aδ-LTMR module neurons.

**Figure 4.**
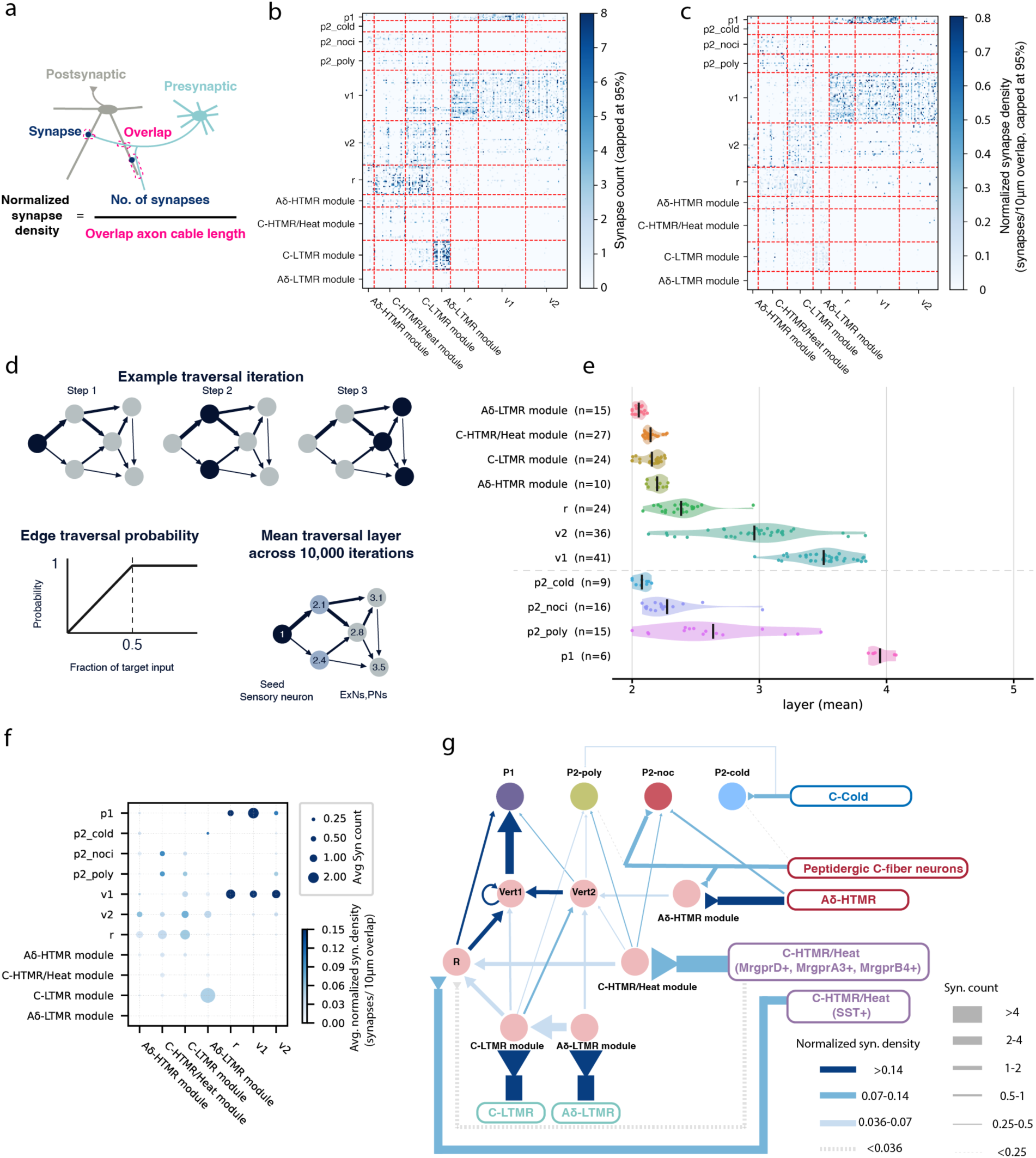
Wiring diagram of sensory processing circuits in the superficial dorsal horn. **a**, Schematic illustrating approach used to quantify (i) axon–dendrite overlap (region for potential synapse formation; axon length within 5 µm of a postsynaptic dendrite) and (ii) normalized synapse density (synapse count normalized by axon–dendrite overlap) to control for spatial proximity between neurons. **b**, Connectivity matrix of synapse counts between individual reconstructed neurons, grouped by subtype (presynaptic neurons, columns; postsynaptic neurons, rows). Values are capped at 8 synapses (95th percentile) for visualization. **c**, Matrix of normalized synapse density between presynaptic and postsynaptic neurons. **d**, Schematic of the probabilistic network traversal (Schlegel et al., 2021) used to assign ExNs and PNs to layers relative to sensory afferents. 135 Sensory afferents (Αβ-LTMRs excluded) were designated as layer 1, and traversal probability was determined by the fraction of input contributed by each presynaptic neuron (threshold, 0.5). Each neuron’s layer was defined as the mean step at which it was first reached by traversal from the seed sensory neurons across 10,000 iterations; neurons reached directly from the sensory seeds therefore occupied earlier layers, whereas those reached through intermediate ExNs occupied later layers. **e**, Probabilistic network traversal layers of ExN and PN subtypes relative to sensory afferents, which were assigned as layer 1. Dots represent individual neurons, violins show their distributions and black bars indicate subtype means. **f**, Summary of subtype-level connectivity. Dot size indicates the mean synapse count per overlapped presynaptic–postsynaptic neuron pair; color indicates mean synapse density, capped at 95% percentile. Only overlapping pairs were considered when mean synapse counts and density were calculated. **g,** Wiring diagram summarizing sensory neuron–ExN–PN connections. Sensory neuron inputs are shown as lines with triangular endpoints, and local spinal connections are shown as arrows. Line thickness denotes mean synapse count per overlapped pairs, and arrow color denotes mean synapse density per overlapped pairs. Only connections with mean synapse count ≥0.25 and synapse density ≥0.036 are shown (solid lines). Three additional sensory inputs that did not meet these thresholds but accounted for more than 3.5% of the total synaptic input to the postsynaptic population are shown as dashed lines: peptidergic C-fiber neuron input to P2-Polymodal PNs; peptidergic C-fiber neuron input to P2-Cold PNs; and C-HTMR/Heat (MrgprD^+^, MrgprB4^+^ *and* MrgprA3^+^) neuron input to radial cells.

To begin to assess the flow of sensory signals from somatosensory neurons to PNs, we applied a probabilistic graph-traversal model (Dorkenwald et al., 2024; Schlegel et al., 2021) to the network of reconstructed sensory neurons, ExNs, and PNs, using the 135 sensory neurons as seeds assigned to layer 1. At each step, a neuron had some probability of traversing to each of its postsynaptic targets; a target neuron was designated as traversed if this probabilistic transition succeeded. The transition probability for a given target increased with the fraction of its total reconstructed-network synaptic input contributed by the presynaptic neuron, saturating at 100% once that fraction reached 50%. We repeated the traversal 10,000 times and defined each neuron’s layer as the mean step at which it was first traversed (**Figure 4d,e; Figure S7h**). Thus, a neuron’s layer represents its average network distance from primary sensory neurons: neurons receiving stronger direct sensory input are reached earlier, and exhibit a low layer value, whereas neurons reached primarily through intermediate ExNs occupy later layers (higher layer values). This analysis revealed a feedforward hierarchy in which the different ExN and PN subtypes occupied distinct network positions. ExNs in the Aδ-HTMR, C-HTMR/Heat, C-LTMR and Aδ-LTMR sensory modules were all positioned near the second layer (**Figure 4e**), indicating that they are mainly driven by direct sensory input of a single modality. Indeed, these four sensory module ExNs receive dense monosynaptic sensory input but little input from other ExNs (**Figure 4b**). This limited ExN input to sensory modules did not reflect a lack of opportunity for synapse formation: despite extensive axon–dendrite overlap (**Figure S7a**), inter- and intra-sensory module connectivity remained sparse, with connection probabilities below 16% and fewer than 0.36 synapses per overlapping neuron pair (**yellow region in Figure S7b,c**). One exception to this general rule was observed for connections between ExNs of the Aδ-LTMR module and the C-LTMR module, which exhibited a relatively high number of synaptic connections, revealing a site of convergence between fast- and slow-conducting LTMR pathways (**Figure 4b**). These findings indicate that the sensory modules function primarily as first-order recipients of modality-specific sensory input and relay this information to downstream spinal cord neuron subtypes.

Beyond these first-order sensory modules, the remaining ExN subtypes occupied deeper positions in the traversal hierarchy. Radial cells received direct Sst^+^ C-HTMR/Heat neuron input (**Figure 2b; Figure S7d**) and occupied a position between the second and third network layers.

Vertical cells received little monosynaptic sensory input and were assigned predominantly to the third network layer (**Figure 4e**). These later layer positions indicate that radial and vertical cells are recruited largely through polysynaptic pathways involving upstream ExNs. Surprisingly, the four lamina I PN subtypes occupied markedly different positions in the traversal hierarchy (**Figure 4e**). P2-Cold and P2-Nociceptive PNs were positioned near the second layer of the traversal model, like the four sensory modules, indicating that their synaptic inputs are dominated by primary sensory neurons. In stark contrast, P2-Polymodal PNs and P1 PNs occupied later network layers, with P1 positioned close to the fourth layer, the deepest layer of the network.

To visualize how these ExN and PN layer positions arise from the underlying connectivity, we next constructed a wiring diagram linking sensory neuron subtypes and the synaptically defined ExN and PN subtypes after excluding weak connections from the subtype-level connectivity matrices (**Figure 4f,g**). This wiring diagram showed that the four PN subtypes are embedded in distinct subnetworks that integrate different sensory modalities through specialized circuit architectures.

The first major pathway linked LTMR, C-HTMR/Heat, and, to a lesser extent, Aδ-HTMR inputs to P1 PNs through a multilayered feedforward network (**Figure 4g**). This network included all sensory modules, radial cells, and both Vert neuron subtypes. Within the P1 PN network, radial and Vert2 neurons consistently showed the highest numbers of synapses and connection probabilities among the downstream ExN targets of all four sensory modules (**Figure 4f; Figure S7b,c**). Both radial and Vert2 neurons, in turn, connected strongly to Vert1 neurons, with connection probabilities of 60% and 42% and mean synapse numbers of 1.51 and 1.21 per overlapping pair, respectively (**Figure S7b,c**). Strikingly, Vert1 neurons uniquely exhibited a prominent recurrent network with one another (42.8% connection probability; 0.94 synapses per overlapping pair). Vert1 neurons also uniquely made dense synapses onto the six P1 ALT neurons (72.7% connection probability; 2.01 synapses per overlapping pair). Thus, with minimal monosynaptic sensory input (<3% of total synaptic inputs) and strong input from deep network layer Vert1 neurons, all six P1 PNs were assigned to the fourth layer of the network. This multilayered feedforward pathway to P1 PNs accounted for 64.4% of all synapses in the network, despite representing only 28.0% of total neurite overlap in the analyzed matrices (**Figure 4b; Figure S7a**). Together, these results indicate that P1 PNs receive predominantly LTMR and C-HTMR/Heat information, with more limited Aδ-HTMR input, through a multilayered excitatory pathway involving the four sensory modules, radial cells, and both vertical cell subtypes.

Unlike P1 PNs, which have minimal monosynaptic sensory inputs, P2-Polymodal PNs received modest monosynaptic input from C-Cold thermoreceptors and peptidergic C-fiber neurons (**Figure S7g**). Interestingly, P2-Polymodal PNs also integrated weak inputs from multiple ExN populations (**Figure 4f; Figure S7b,c**), including the C-LTMR module (16% connection probability; 0.36 synapses per overlapping pair), the C-HTMR/Heat module (24%; 0.42 synapses per overlapping pair), and Vert2 neurons (17%; 0.31 synapses per overlapping pair). This combination of direct C-Cold neuron and peptidergic C-fiber neuron inputs and polysynaptic LTMR and C-HTMR/Heat neuron inputs placed P2-Polymodal PNs between the second and third layers in the graph traversal model. Thus, P2-Polymodal neurons broadly integrate tactile, thermal, and nociceptive information through a mixture of monosynaptic and polysynaptic inputs (**Figure 4g**).

P2-Cold and P2-Nociceptive neurons stood out in our analysis because they received a much higher fraction of monosynaptic inputs from C-Cold thermoreceptors and nociceptors, respectively, compared to P1 and P2-Polymodal PNs (**Figure S7g**). Remarkably, both P2-Cold and P2-Nociceptive PNs received little input from lamina I-II ExNs (**Figure 4f**), except for modest C-HTMR/Heat module input to P2-Nociceptive PNs (27% connection probability; 0.44 synapses per overlapping pair). As a result, both of these PN subtypes were assigned close to the second layer of the graph traversal model. Thus, in sharp contrast to the polysynaptic, multilayered lamina I-II ExN pathway converging onto P1 PNs, P2-Cold and P2-Nociceptive PNs predominantly received monosynaptic inputs and are dedicated to conveying cold and nociceptive information, respectively, to the brain (**Figure 4g**).

Altogether, this analysis allowed us to construct a lamina I-II wiring diagram linking sensory neuron, ExN, and PN subtypes (**Figure 4g**). We found that the four connectivity defined lamina I PN classes integrate distinct sensory modalities through markedly different circuit architectures for output to the brain. LTMR and C-HTMR/Heat signals are mainly routed through a multilayered feedforward ExN network involving recurrently connected Vert1 neurons (putative GRPR*^+^* neurons) before converging densely onto P1 PNs (putative Gpr83*^+^* neurons). This architecture is consistent with a previous report that Gpr83*^+^* ALT PNs convey light touch-related signals to the brain (Choi et al., 2020). Moreover, C-HTMR/Heat neurons and GRPR^+^ spinal neurons (Vert1 neurons) play key roles in itch sensation (Huang et al., 2018; Mu et al., 2017; Sheahan et al., 2024; Sun and Chen, 2007). Thus, P1 PNs likely convey innocuous mechanosensory, warm thermal, and pruriceptive information to the brain. P2-Polymodal PNs integrate weaker but broader combinations of direct C-Cold and peptidergic C-fiber neuron inputs together with polysynaptic LTMR and C-HTMR/Heat inputs, thus likely supporting broad, polymodal (tactile, thermal, and nociceptive) tuning. In contrast, P2-Cold and P2-Nociceptive PNs receive predominantly direct monosynaptic input from highly select sensory afferent subtypes, forming ascending pathways for conveying cold and nociceptive information to the brain.

### Feedforward inhibition selectively gates sensory modules

We next asked how inhibitory circuit motifs engage the four synaptically defined laminae I ALT PNs to shape modality-specific processing for output to the brain. To address this, we focused on the subsets of inhibitory neurons (IhNs), characterized in parallel work (Xiang et al., 2026a), that mediate homotypic presynaptic inhibition and provide feedforward inhibition (**Figure S8a,b**).

We refer to these IhN populations as Islet-PSI-IhN^C-HTMR/Heat^, Islet-PSI-IhN^C-LTMR^, Islet-PSI-IhN^Aδ-LTMR^, and PSI-IhN^Aδ-HTMR^ (Xiang et al., 2026a). We estimate that these IhN populations comprise approximately 30% of lamina I-II IhNs since, in the densely reconstructed region, 9 of 31 IhNs both exhibited Islet-like morphology and formed axoaxonic synapses.

We examined the postsynaptic spinal targets of these four lamina I-II IhN subtypes. Mapping their synapses onto the ExN subtypes defined above showed that most of these IhNs targeted their homotypic ExN modules, while a subset also formed a substantial number of heterotypic feedforward inhibitory synapses (**Figure S8c-f**). Islet-PSI-IhN^C-HTMR/Heat^ neurons received inputs exclusively from C-HTMR/Heat neurons and formed the majority of their axodendritic and dendrodendritic synapses onto ExNs within C-HTMR/Heat modules (Davis et al., 2023), thereby mediating homotypic feedforward inhibition. Similarly, Islet-PSI-IhN ^Aδ-LTMR^ provided homotypic feedforward inhibition by forming axodendritic synapses onto Aδ-LTMR modules. In contrast, Islet-PSI-IhN ^C-LTMR^ and PSI-IhN^Aδ-HTMR^s, in addition to targeting their homotypic sensory pathways, formed a substantial number of dendrodendritic and axodendritic synapses onto C-HTMR/Heat modules, thereby mediating heterotypic feedforward inhibition. This pattern is consistent with our parallel work reporting heterotypic FFI motifs are biased toward spinal neurons that receive C-HTMR/Heat inputs (Xiang et al., 2026a). This result also demonstrates that a subset of these heterotypic FFI connections directly target excitatory neurons within the C-HTMR/Heat modules. Notably, PSI-IhN^Aδ-HTMR^s also directly inhibited P2-Nociceptive PNs, consistent with a role in suppressing nociceptive output to the brain.

Together, these findings show that all sensory modules, as well as the P2 nociceptive PN population that densely sample select sensory inputs, are subject to homotypic feedforward inhibition (**Figure S8g**). This inhibitory motif may moderate the feedforward excitatory drive within individual sensory channels. Interestingly, LTMR and Aδ-HTMR nociceptor pathways also engage heterotypic feedforward inhibition of C-HTMR/Heat module neurons. This organization presumably enables cross-modality interactions and may sharpen force tuning acuity (See Xiang et al., 2026a).

### A connectome-based model predicts PN sensory selectivity

Thus far, we have established a wiring diagram of the superficial dorsal horn microcircuitry and identified modality-specific circuit architectures for outputs to the brain. We next asked how such connectivity might shape the transformation and routing of sensory signals to the different PN output pathways. To address this, we built a firing rate-based model parameterized by the measured synaptic connectivity. In this model, sensory neurons provide feedforward excitation to reconstructed ExNs, IhNs, and PNs (**Figure 5a; Figure S9a**).

**Figure 5.**
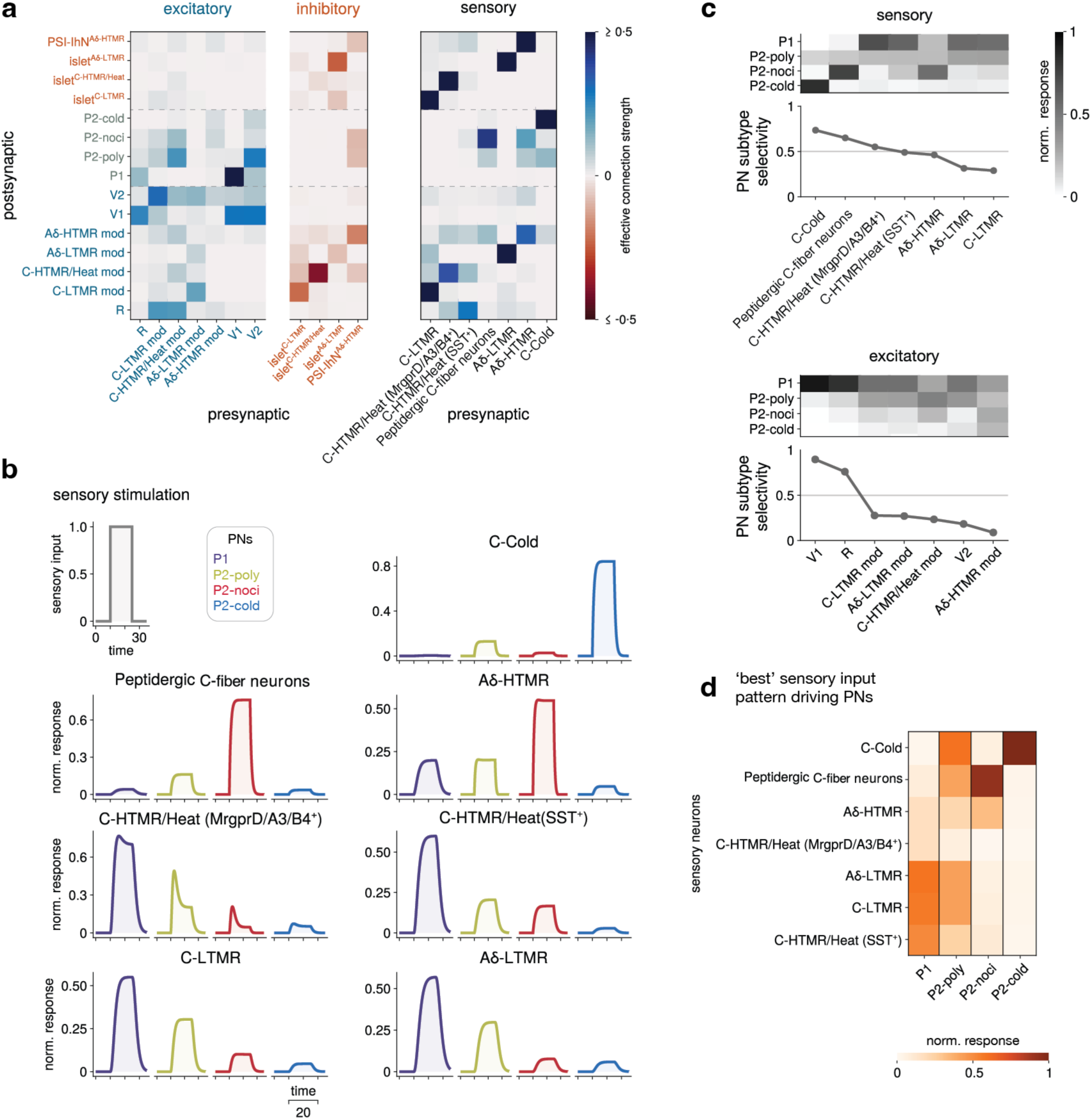
Connectome-constrained modeling of the superficial dorsal horn microcircuit. **a**, Connectivity matrices derived from synapse count data (normalized and scaled; see Methods). The weights in these matrices represent effective synaptic strengths between dorsal horn ExNs, IhNs, and PN subtypes (left; recurrent connectivity) and between sensory neuron subtypes and dorsal horn ExNs (right; feedforward connectivity). For a schematic wiring diagram based on these matrices, see **Figure S9a**. **b**, Simulated PN responses to stimulation of individual sensory neuron subtypes. In each condition, responses are normalized by the sum of the steady-state values across the four PN subtypes. **c**, Top: heatmap of normalized PN responses across all sensory neuron subtypes (upper panel) and the corresponding PN subtype selectivity index (lower panel). The selectivity index equals 1 when only a single PN subtype is activated and 0 when the two most-activated PN subtypes are equally activated (see Methods). Sensory neurons are ordered by this selectivity index value. Bottom: same analysis repeated for stimulation of ExN subtypes. For analysis of the sensitivity of these results to model parameters, i.e., the strengths of inhibition and excitation, see **Figure S9b-d** and Methods. **d**, Heatmap of optimal sensory neuron inputs that maximally drive each PN subtype. Each column corresponds to the model input that elicits the largest response for a given PN subtype, subject to unit-norm and non-negative input constraints (see Methods).

We first used this model to predict responses of the four PN subtypes to stimulation of individual sensory afferent classes (**Figure 5b**). We compared steady-state PN responses across sensory stimulus conditions and, using a selectivity metric, quantified the degree to which each sensory neuron subtype drives each of the four connectivity defined PN subtypes (**Figure 5c**; see Methods). As predicted, stimulation of the different sensory neuron subtypes elicited distinct PN response patterns. C-cold neurons and peptidergic C-fiber neurons selectively activated P2-Cold and P2-Nociceptive PNs, respectively, with only weak responses observed in the other PN subtypes. In contrast, C-LTMR and Aδ-LTMR stimulation activated PNs in a more distributed manner, producing the strongest response in P1 PNs, an intermediate level response in P2-Polymodal PNs, and very weak responses in P2-Nociceptive and P2-Cold PNs. To assess the contribution of excitatory interneurons independent of sensory input, we performed simulations with direct stimulation of ExNs and found that Vert1 cells and radial cells exhibited the largest degree of selectivity, predominantly activating P1 PNs (**Figure 5c**).

Finally, our simulations revealed sensory input patterns that maximally drive each PN subtype (**Figure 5d**; see Methods). Complementing the individual sensory neuron activation simulations above, this analysis revealed that different PN subtypes are preferentially driven by distinct sensory input combinations: P2-cold PNs are mainly driven by C-Cold afferents; P2-nociceptive PNs by combined peptidergic C-fiber neuron and Aδ-HTMR nociceptor inputs; P1 PNs by C-LTMR, Aδ-LTMR, and C-HTMR/Heat inputs; and P2-polymodal PNs by a broad combination of LTMR, C-HTMR/Heat, C- and Aδ-nociceptors, as well as C-Cold neuron inputs (**Figure 5d**). Together, these simulations suggest that signals representing cold, noxious thermal, and noxious mechanical stimuli are transmitted through relatively selective lamina I PN output pathways, whereas light and medium force tactile signals as well as pruriceptive signals are transmitted via P1 PNs and to a lesser extent two additional PN subtypes. These findings are consistent with modality-specific input routing predicted by the wiring diagram (**Figure 4e**) and available *ex vivo* slice electrophysiology measurements (Bohic et al., 2026; Choi et al., 2020; Lezgiyeva et al., 2026; Razlan et al., 2025; Zhang et al., 2024).

## Discussion

Here, we used cell type-specific genetic labeling (Zhang et al., 2019) in conjunction with synapse-resolution connectomics (Phelps et al., 2021) to ask how sensory neuron signals are integrated in the superficial dorsal horn for output to the brain via lamina I ALT PNs. Our findings support a modular organization linking sensory afferents to four connectivity-defined classes of lamina I PNs. This segregation of synaptically-defined ALT output channels provides a synaptic explanation for classic physiological recordings in primates and cats describing cooling-selective, nociceptive-selective, and wide-dynamic-range lamina I PNs (Christensen and Perl, 1970; Han et al., 1998), and aligns with recent mouse studies identifying genetically defined PN subpopulations with distinct properties and functions (Choi et al., 2020; Razlan et al., 2025; Sheahan et al., 2024; Warwick et al., 2022). The P2-Cold PN population described here likely corresponds to cold-selective PNs. These PNs have dorsally restricted dendritic arbors and receive predominantly C-Cold sensory neuron inputs and few excitatory inputs from lamina I-II ExNs. These synaptic and morphological features also align with the reported properties of cold-selective PNs labeled by Calbindin^+^ neurons of the Phox2a^+^ lineage of ALT PNs (Razlan et al., 2025; Roome et al., 2020). A second class, P2-Nociceptive PNs, receives near-exclusive direct sensory input from Aδ-HTMRs (mechano-nociceptors) and peptidergic C-fiber neurons, including C-Heat (Sstr2^+^) nociceptors, consistent with nociceptive-selective PNs. A third PN population, P2-Polymodal PNs, broadly integrates weak monosynaptic C-Cold neuron and peptidergic C-fiber neuron inputs with weak polysynaptic C-LTMR- and C-MH-module ExN input, a convergence that likely underlies the polymodal nature of wide dynamic range PNs. The dorsally restricted dendritic arbors and direct nociceptor drive of P2-Nociceptive and P2-Polymodal PN populations are consistent with the morphology and synaptic input profile of Tacr1^+^ PNs (Choi et al., 2020). Finally, P1 PNs resemble Gpr83^+^ PNs in both morphology and their sensory input profile, extending ventrally directed dendrites and receiving strong, mostly polysynaptic drive from LTMRs and several C-fiber mechanosensory neuron subtypes (Choi et al., 2020). P1 PNs may also correspond to “giant cells” described in previous anatomical studies, a largely Tacr1 negative population of lamina I PNs that similarly receives little direct primary afferent input but extensive excitatory input from dorsal horn interneurons (Cameron et al., 2015; Polgar et al., 2008; Puskar et al., 2001). Thus, P1 PNs represent the principal lamina I ALT output pathway engaged by vertical cells and the extensive polysynaptic, feedforward excitatory dorsal horn circuitry within lamina I-II. Together, these findings reveal a circuit logic of the superficial dorsal horn where modality specific DRG sensory neuron signals are differentially routed via specific circuit motifs to one or more discrete lamina I PN classes.

The contrast between the monosynaptic relay of cold and nociceptive signals (conveyed by C-Cold neurons, and C-peptidergic neurons and Aδ-HTMRs, respectively) and the polysynaptic routing of light and medium force tactile signals (conveyed by LTMRs and C-HTMR/Heat neuron populations) may reflect the distinct functional demands of these sensory channels. Activation of nociceptors such as Aδ-HTMRs and Sstr2^+^ C-Heat neurons, as well as their downstream Tacr1+ PNs, elicits robust nocifensive behaviors and aversion (Arcourt et al., 2017; Choi et al., 2020; Ghitani et al., 2017; Lezgiyeva et al., 2026). By contrast, LTMR and C-HTMR/Heat afferents mediate a broader repertoire of functions and behaviors. Indeed, C-LTMRs and the four C-HTMR/Heat afferent classes (Sst^+^, MrgprD^+^, MrgprA3^+^, and MrgprB4^+^), or combinations or subsets of them, are implicated in pain (Dong et al., 2001; Sharif et al., 2020), chemical itch (Han et al., 2013; Liu et al., 2009; Solinski et al., 2019), mechanical itch (Hill et al., 2022), tickle (Zotterman, 1939), wet dog shakes (Zhang et al., 2024), social touch (Elias et al., 2023; Liu et al., 2025), and pleasant touch (Loken et al., 2009; Vrontou et al., 2013). Despite this functional diversity, LTMR and the four C-HTMR/Heat neuron subtype signals are mainly routed through the elaborate vertical cell feedforward network to target P1 PNs. Such a complex, polysynaptic architecture enables greater integrative and coding capacity than the direct monosynaptic relay organization observed in the P2-Cold and P2-nociceptive PN pathways. The much weaker monosynaptic sensory neuron input to P1 PNs, relative to the P2 PN subtypes, further suggests that the vertical cell–P1 output pathway is more strongly shaped by CNS inputs, including descending signals that may reflect different internal states (Liu et al., 2019; Nguyen et al., 2023). We propose that the sensory neuron–sensory module–radial cell/Vert2–Vert1–P1 PN architecture, together with recurrent connections among Vert1 neurons, provides a substrate for population coding and internal-state-dependent modulation of light- and medium-force tactile, warm thermal, and pruriceptive signals before their transmission to the brain via the ALT. Consistent with this idea, activation of Gpr83^+^ PNs, which we believe are P1 neurons, can generate either appetitive or aversive valence depending on their pattern of activation (Choi et al., 2020). Similarly, GRPR^+^ vertical cells have been implicated in both itch and pain (Polgar et al., 2023). Future studies defining CNS as well as polysynaptic Aβ-LTMR subtype inputs to the vertical cell–P1 PN pathway, as well as the physiological responses of the four synaptically defined lamina I PNs under naturalistic stimulation conditions in awake animals, will inform our understanding of how diverse sensory inputs are transformed into spinal output signals underlying distinct reflexes and percepts.

Larger scale circuit reconstructions and identification of additional physiologically distinct sensory channels will be required to achieve a full appreciation of the flow of signals from sensory neurons to ALT output neurons. Many reconstructed neurons extend beyond the EM volume and likely form additional synapses outside the dataset. For instance, vertical cells can synapse onto neurons in the lateral spinal nucleus, which houses ALT PNs (Polgar et al., 2023; Quillet et al., 2023); these longer-range vertical cell connections are difficult to capture with current connectomic methods. Likewise, the reconstructed circuits described here almost certainly receive CNS inputs that arise from neurons whose somata lie outside the EM volume and cannot currently be identified. Additional limitations of the present study include our inability to distinguish between the synapses of MrgprD^+^, MrgprA3^+^ and MrgprB4^+^ C-HTMR/Heat neuron subtypes. Similarly, while we grouped together all CGRP^+^ C-fiber neurons in the present analysis, the physiological properties of some transcriptionally defined CGRP^+^ C-fiber neuron subtypes are not fully appreciated (Qi et al., 2024), nor can we currently resolve their lamina I-II synapses. Despite these limitations, the reconstructed circuitry presented here provides an organizational logic of the core superficial dorsal horn circuitry. Moreover, the dSC1 EM volume and SPINE analysis will enable connectomic reconstruction of lamina III and IV neurons. This includes deep dorsal horn ALT PN populations (antenna cells) (Bell et al., 2024; Schoenen, 1982; Todd, 1989) as well as postsynaptic dorsal column pathway neurons. The latter form a major spinal ascending pathway for Aβ-LTMR inputs that were not characterized in the current study (Abraira and Ginty, 2013). In addition, broadening coverage of inhibitory dorsal horn neurons beyond those mediating presynaptic inhibition (∼30% of the lamina I/II IhNs, studied here) will reveal the full extent of inhibitory circuit motifs governing superficial and deep PN output pathways.

By leveraging genetic labeling and neuronal morphology, we were able to infer the identities of some PNs and the two vertical cells subtypes, thereby linking the lamina I-II circuit map described here to prior molecular, physiological, functional, and anatomical insights. Looking forward, it will be useful to establish genetic access of connectivity-defined spinal cord neuron subtypes identified in the present study to enable physiological analyses and targeted functional perturbations of circuit nodes for testing hypotheses derived from the connectome. More broadly, as connectomic approaches become more routine, bidirectional mapping between connectivity-, physiologically-, and molecularly defined cell types, during development, in other animals including humans, and in disease states will help achieve a deeper understanding of the organizational logic of the somatosensory system.

**Figure S1.**
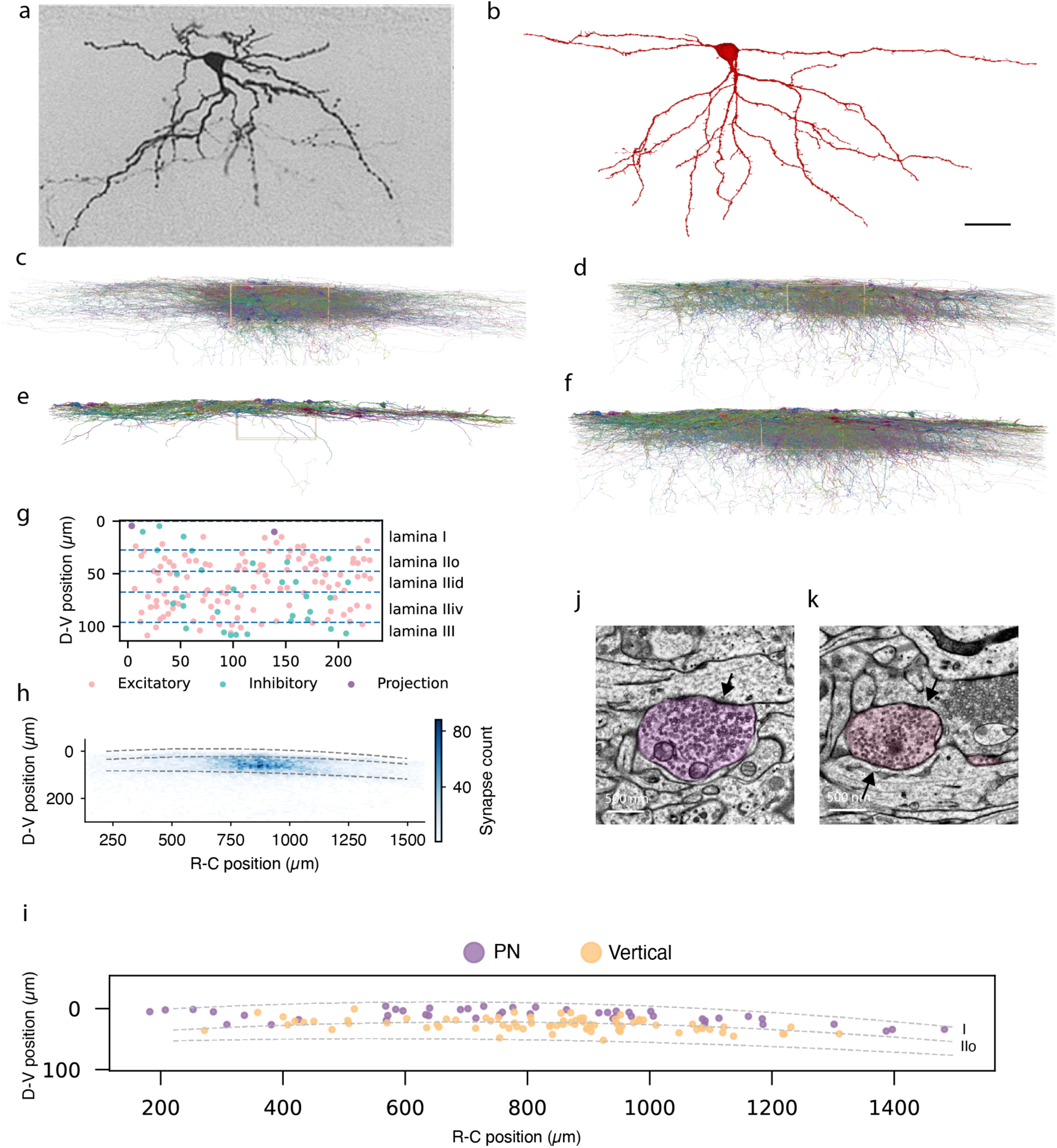
Dense reconstruction of ExNs. **a**, Classic vertical cell labeled by biocytin filling. Figure adapted from (Grudt and Perl, 2002). **b**, A vertical cell identified in the EM volume. Vertical cells can be reliably recognized by their characteristic laminar location and ventrally directed dendrites. Scale bar, 30 µm. **c**, Dense reconstruction of excitatory neurons (*n* = 100) within the ROI indicated by the yellow box, which spans 230 × 100 × 31 µm along the rostrocaudal, dorsoventral, and mediolateral axes, respectively. **d**, Reconstructed vertical cells (n = 77) shown relative to the ROI. **e**, Reconstructed lamina I projection neurons (n = 46) shown relative to the ROI. **f**, All reconstructed neurons included in this study (n = 223) shown relative to the ROI. **g**, Soma distribution of ExNs, IhNs, and PNs within the reconstruction ROI. **h,** Spatial distribution of output synapses from reconstructed ExNs in the dSC1 volume. Color intensity represents synapse density. Dashed lines indicate the boundaries between white matter and lamina I, lamina I and lamina II, and lamina II and lamina III. Only the superficial 300 µm region below the white matter–lamina I boundary is shown. **i**, Soma distribution of all reconstructed vertical cells and PNs across the EM volume. These cell types were reconstructed throughout the volume to better capture sensory processing circuitry. **j**, Example excitatory synapse. Excitatory neurons in laminae I-II form classic asymmetric synapses with a prominent postsynaptic density (arrow). **k**, Example inhibitory synapse. Inhibitory neurons in laminae I-II form classic symmetric synapses with little or no postsynaptic density (arrow). This neuron also forms an axoaxonic synapse onto a sensory terminal, further supporting its inhibitory identity.

**Figure S2.**
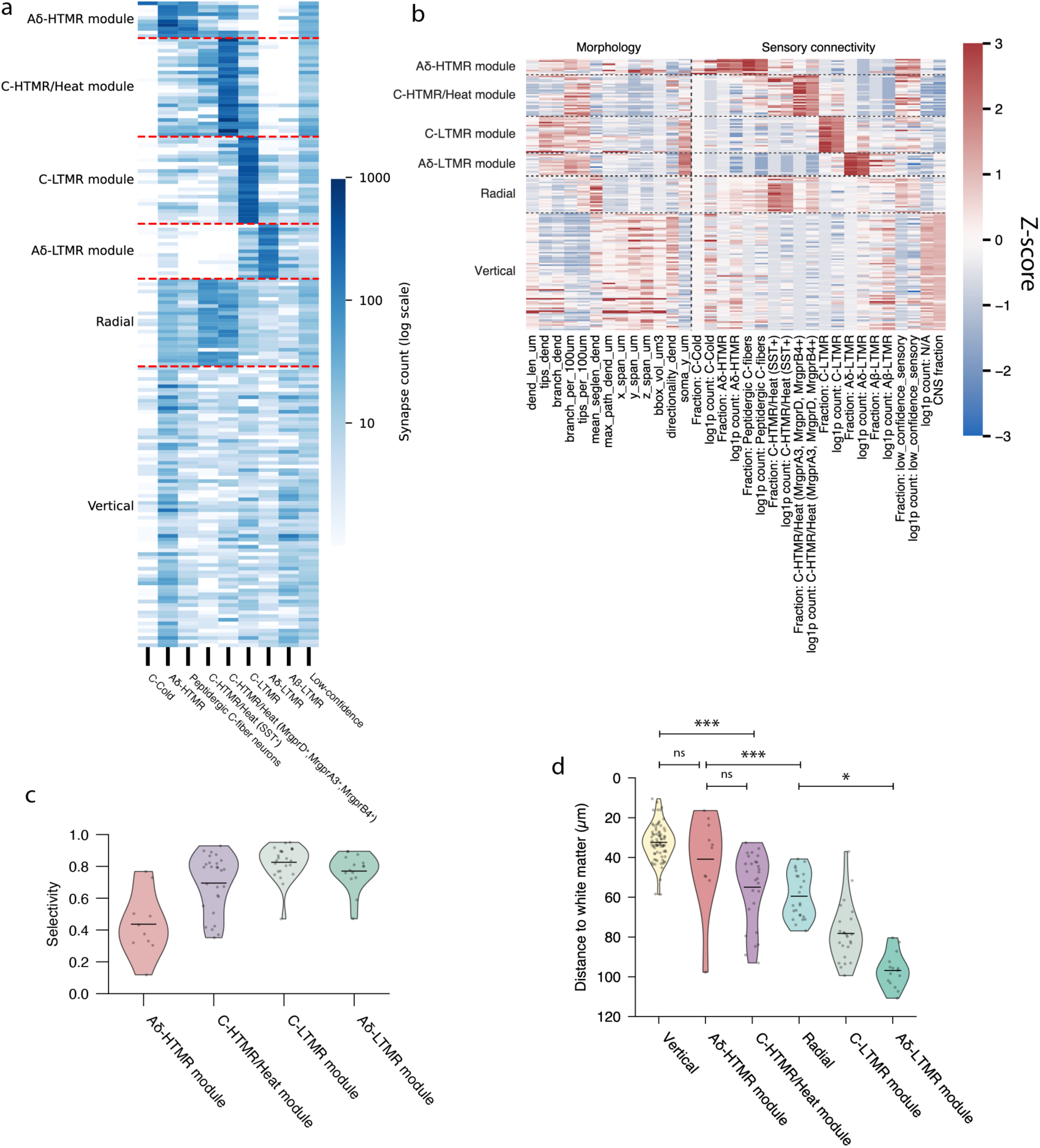
ExN clustering. **a**, Raw sensory neuron synapse counts onto individual ExNs. Rows show single ExNs sorted by subtype; columns indicate sensory neuron subtypes; color denotes synapse count. **b**, Z-score heatmap of the morphological and synaptic input features used for UMAP embedding. ExNs within the same subtype show similar feature profiles; features are described in Methods. **c**, Selectivity of sensory input across ExN modules. Violin plots show, for each sensory-defined ExN module, the fraction of total sensory synaptic input contributed by the module’s defining sensory subtype (dots, individual ExNs; line, median). **d**, Depth of ExN somata measured as distance from the white matter boundary. *P<0.05;**P<0.01;***P<0.001, using a Kruskal–Wallis test, and Dunn posthoc test with Holm– Bonferroni-correction. Vertical cells, n=77; radial cells, n=24; Aδ-LTMR module ExNs, n=15; C-LTMR module ExNs, n=24; Aδ-HTMR module ExNs, n=10; C-HTMR/Heat module ExNs, n=27.)

**Figure S3.**
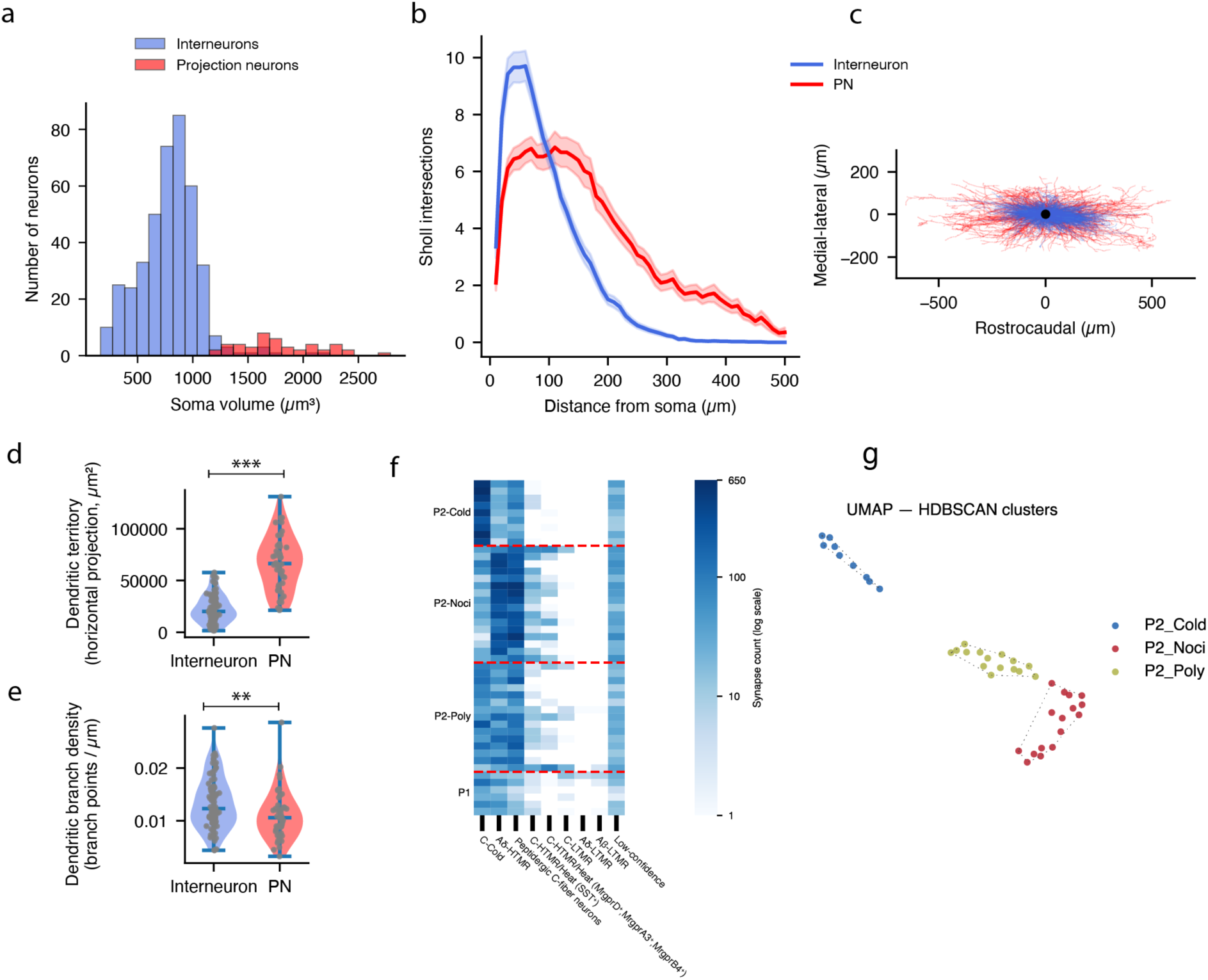
PN morphologies and clustering. **a**, Soma volume distribution of lamina I neurons. Putative projection neurons (n = 45) have a mean soma size 1800.9 ± 365.5 µm^3^ (mean ± SD), lamina I interneurons (n = 411) have mean soma size 787.3 ± 269.0 µm^3^ (mean ± SD). One additional PN, located at the edge of the EM volume, had a partially reconstructed soma and was therefore excluded from soma-volume quantification; inspection of its cross-sections nevertheless confirmed its large soma size. **b**, Sholl analysis of the dendritic arborizations of 46 lamina I PNs and 85 interneurons. PNs displayed complex dendritic branching extending over a larger spatial range. For **Figures S3b-e**, we included in the analyses 85 well-reconstructed interneurons where their dendritic and axonal compartments could be automatically split. **c,** Dendritic fields aligned to the soma. Individual dendritic arbors from PNs (red, n = 46) and interneurons (royal blue, n = 85) were centered on the soma position to compare their spatial organization. PNs displayed broader rostrocaudal dendritic territories, whereas interneurons showed more compact dendritic fields. **d,** Quantification of dendritic territory in the horizontal view as **c**. PNs exhibited significantly larger dendritic fields than lamina I interneurons, measured by the area of the convex hull encompassing dendritic processes in the horizontal projection (PN: mean 67,030.8 ± 26,362.6 µm²; interneurons: mean = 22,724.8 ± 13,471.0 µm²; *p* = 2.2 × 10⁻¹⁶, n_PNs_ = 46, n_interneurons_ = 85, Mann–Whitney U test). **e**, Quantification of dendritic branching density. Interneurons exhibited slightly higher dendritic branch density compared with PNs. Branch density was calculated as the number of branch points divided by total dendritic length (PN: mean 0.011± 0.005 branch points per μm vs. interneuron: mean 0.013 ± 0.005; *p* = 0.0076, n_PNs_ = 46, n_interneurons_ = 85, Mann–Whitney U test). **f**, Raw sensory synapse counts onto individual PNs. Rows show single PNs sorted by subtype. Columns indicate sensory afferent subtypes; color denotes synapse count. **g**, UMAP embedding of PNs using synaptic input features. Dashed outlines indicate HDBSCAN-identified clusters; colors denote the final subtype assignments.

**Figure S4.**
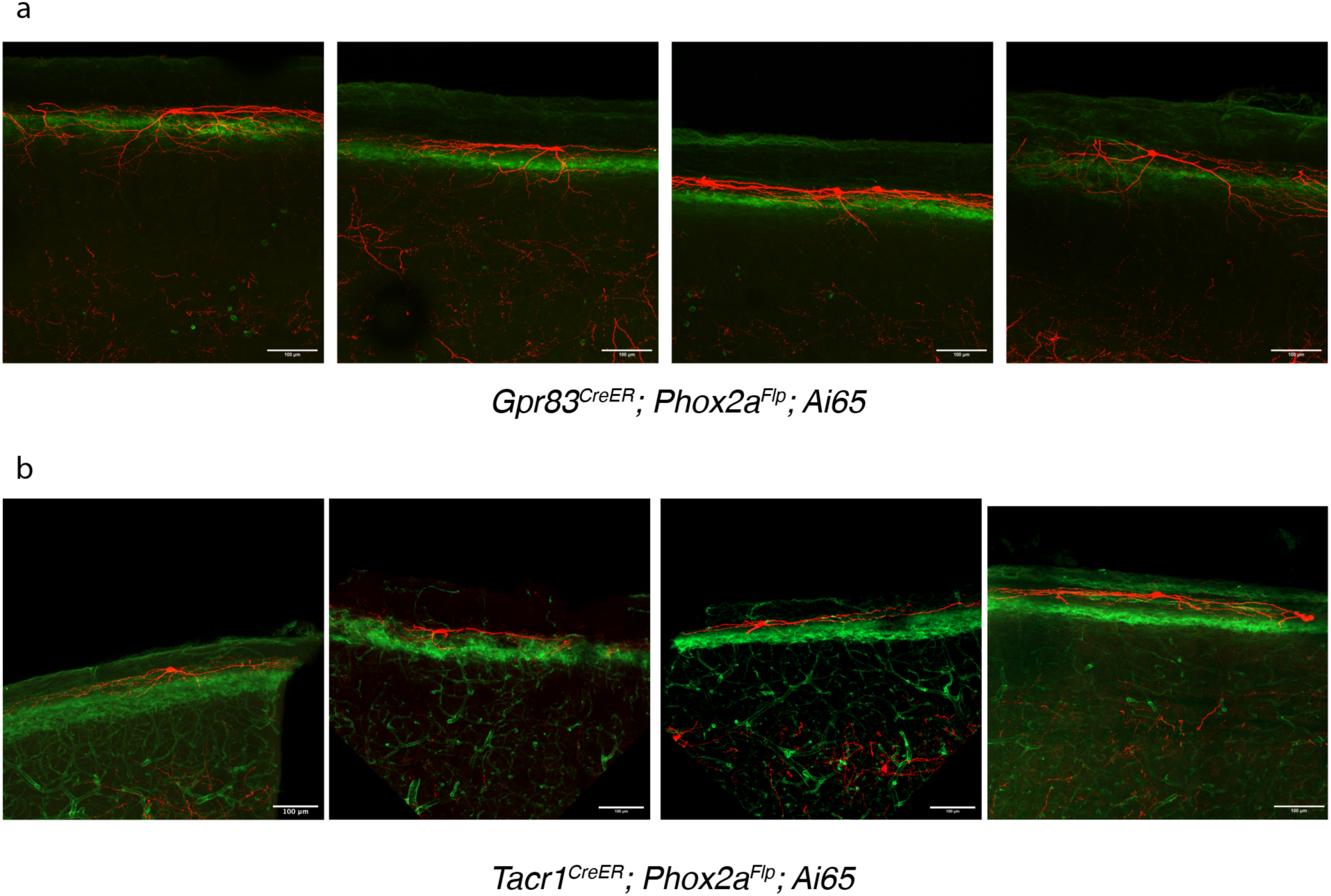
Sparse labeling of *Gpr83^+^* and *Tacr1^+^* lamina I projection neurons. **a**, Confocal microscopy of *Gpr83*⁺ lamina I PN morphology. *Gpr83*⁺ projection neurons (red) extend dendrites ventrally, below the lamina IIid boundary marked by IB4 staining (green). **b**, Confocal microscopy of *Tacr1*⁺ lamina I PN morphology. *Tacr1*⁺ projection neurons (red) extend dendrites that are largely confined to lamina I and dorsal lamina IIo, remaining above the lamina IIid boundary marked by IB4 staining (green). Scale bar, 100 µm (a,b).

**Figure S5.**
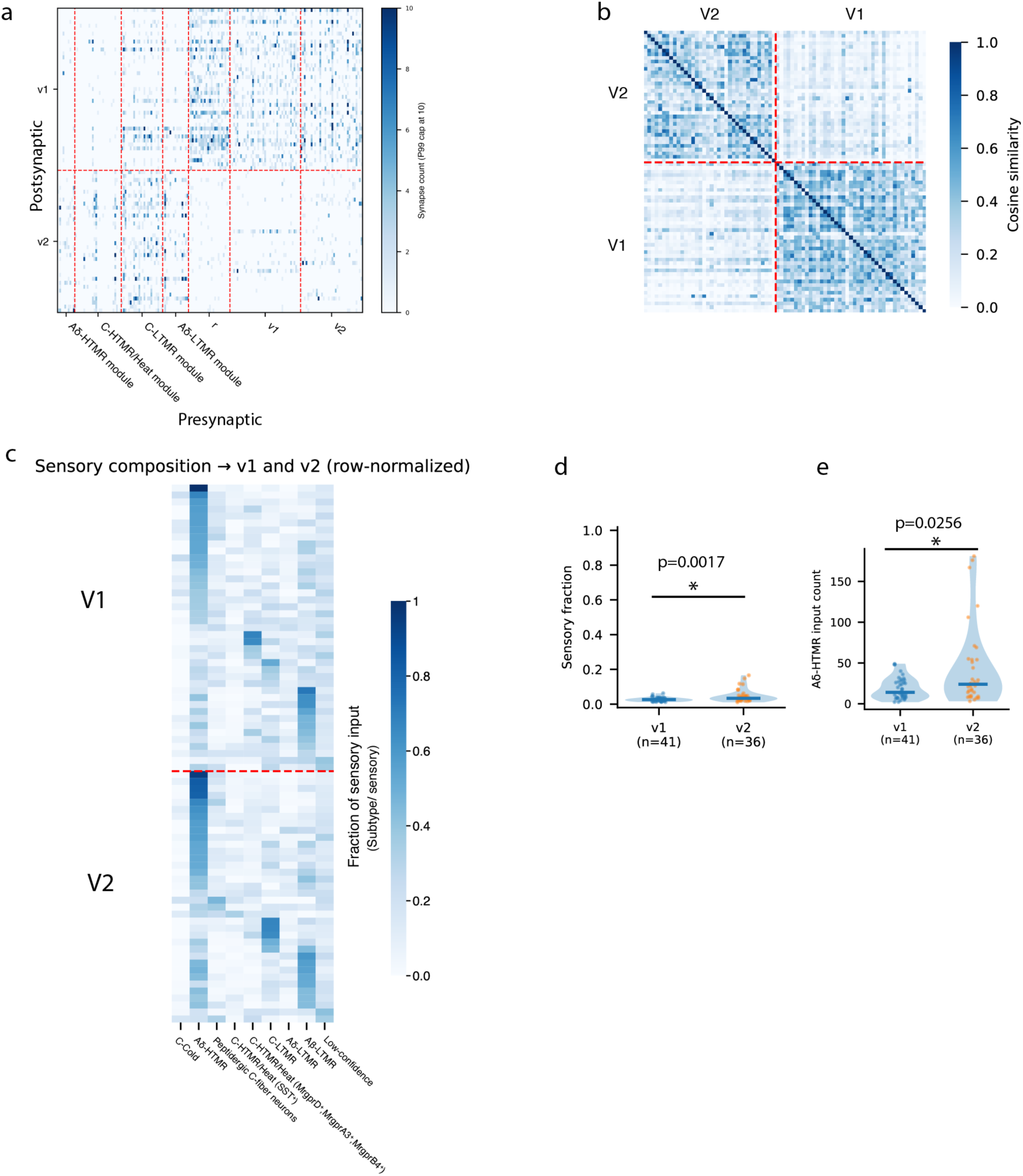
Vert1 neuron and Vert2 neuron connectivity. **a**, ExN inputs onto the two vertical cell subpopulations. Vert1 neurons receive input from the C-LTMR module, radial cells, Vert2 cells, and recurrent input from other Vert1 neurons, whereas Vert2 neurons primarily receive input from the sensory modules. **b**, Cosine similarity matrix of vertical-cell ExN input profiles. Vertical cells segregate into two populations (Vert1 and Vert2) defined by spectral clustering of ExN input features. **c**, Fraction of sensory inputs from each afferent subtype onto Vert1 and Vert2. **d**, Fraction of sensory neuron versus non-sensory neuron synaptic input onto Vert1 and Vert2 neurons. Vert2 neurons receive slightly, but significantly more direct sensory input (p = 0.0017, Mann–Whitney test, v1:n=41, v2: n=36). Note that both Vert1 and Vert2 neurons receive very few direct sensory neuron synaptic inputs. **e**, Raw Aδ-HTMR synapse counts onto Vert1 and Vert2. Vert2 neurons receive slightly, but significantly more Aδ-HTMR inputs compared to Vert1 neurons (p = 0.0256, Mann–Whitney test, v1:n=41, v2: n=36).

**Figure S6.**
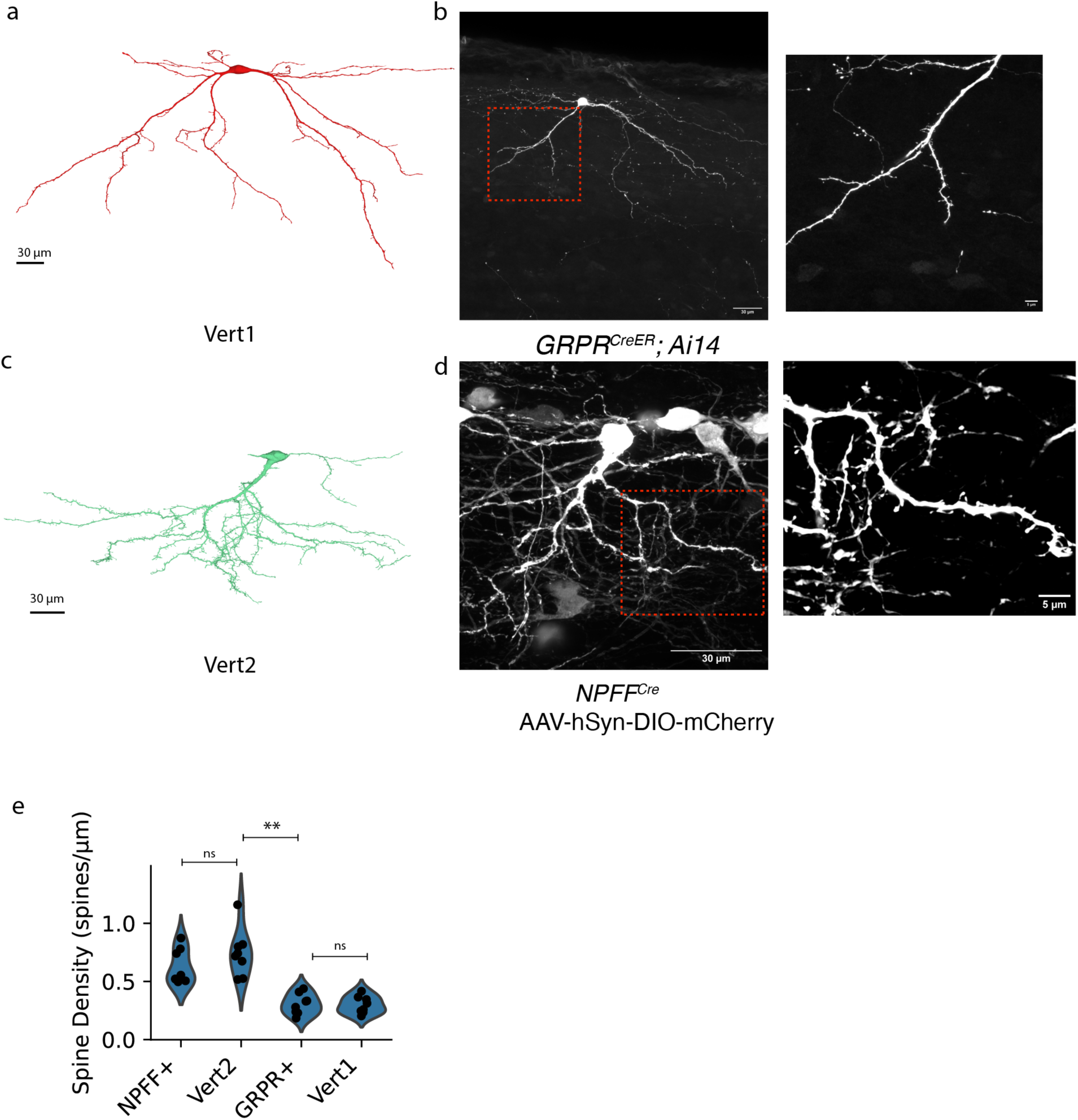
Genetically labelled Grpr^+^ and NPFF^+^ vertical cells visualized by light microscopy match with Vert1 and Vert2 neuron morphologies in the EM dataset. **a**, Example Vert1 neuron morphology reconstructed in the EM volume. **b**, A sparsely labeled GRPR⁺ vertical cell from a *Grpr^CreER^*; *Ai14* mouse. Right, higher-magnification view highlights sparse dendritic spines. **c**, Example Vert2 neuron morphology reconstructed in the EM volume. **d**, A sparsely labeled NPFF⁺ vertical cell (from an *NPFF^Cre^*; AAV-hSyn-DIO-mCherry labeled mouse). Right, higher-magnification view highlights abundant dendritic spines. **e**, Dendritic spine density comparison across light microscopy–labeled neurons and EM-reconstructed vertical cells. NPFF⁺ vertical cells exhibit spine densities similar to Vert2 neurons and significantly higher than GRPR⁺ vertical cells and Vert1 neurons. (p = 2.91 × 10^−5^; NPFF⁺ vs Vert2, p = 1; Vert2 vs GRPR⁺, p = 1.1 × 10^−3^; GRPR⁺ vs Vert1, p = 1; NPFF⁺ vs GRPR⁺, p = 1.2 × 10^−2^, using a Kruskal–Wallis test, and a Dunn’s posthoc with Bonferroni correction. NPFF^+^ cell, n=7; GRPR^+^ cell, n=8; V1 cell, n=8; V2 cell, n=8)

**Figure S7.**
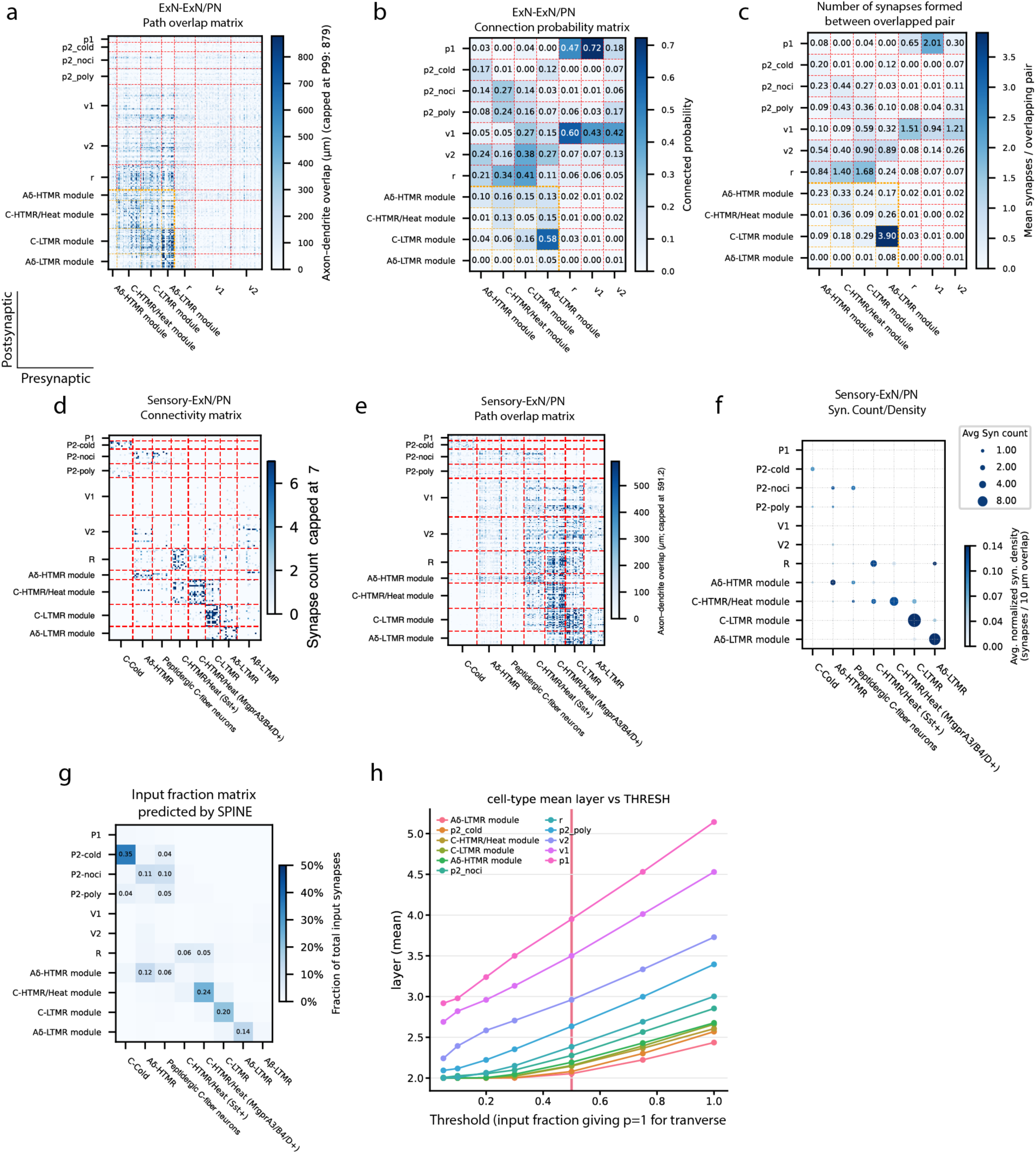
Connectivity and axo-dendritic overlap matrices between sensory neurons, ExN and PN subtypes. **a,** Matrix of axon–dendrite overlap for each ExN–PN pair, quantified as the total presynaptic axon cable length within 5 μm of the postsynaptic dendrite. The sensory module-to-sensory module region is delineated by yellow lines. **b,** Matrix of connection probability, defined as the fraction of ExN–PN pairs with axon–dendrite overlap that formed one or more synapses. The sensory module-to-sensory module region is delineated by yellow lines. **c,** Matrix of mean synapses count per overlapped neuron pair. The sensory module-to-sensory module region is delineated by the yellow lines. **d,** Connectivity matrix summarizing synaptic connections among reconstructed sensory neurons, ExNs, and PNs. **e,** Matrix of axon–dendrite overlap between reconstructed sensory neurons and postsynaptic spinal cord neurons, quantified as in **d**. **f,** Mean synapse number and synapse density for sensory neuron-ExN/PN pairs with axon– dendrite overlap. Synapse density is calculated as the number of synapses per unit overlap length. **g,** Fraction of total synaptic input contributed by each sensory subtype to reconstructed postsynaptic neurons estimated by SPINE. Only connections contributing, on average, more than 3.5% of the total synaptic input to the target ExN/PN subtypes are shown. **h**, Robustness of traversal layers to threshold choice. Mean traversal layer of each ExN and PN subtype across the tested thresholds (2,000 iterations per threshold). Threshold choice altered the absolute layer values but not the relative ordering of ExN and PN subtypes.

**Figure S8.**
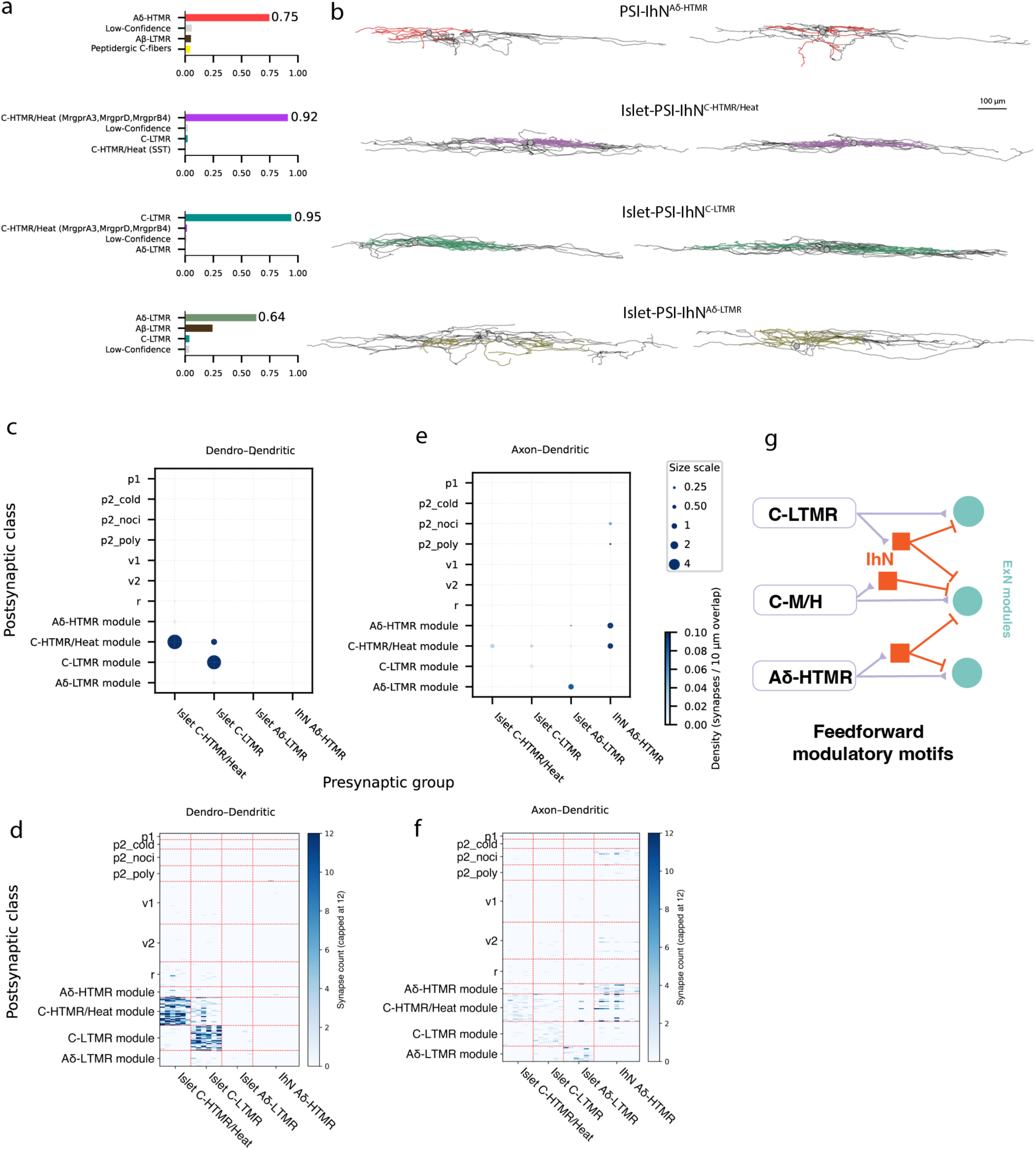
Connectivity between IhNs and ExNs. **a**, Sensory input composition of PSI-IhN^Aδ-HTMR^, islet-PSI-IhN^C-HTMR/Heat^, islet-PSI-IhN^C-LTMR^, and islet-PSI-IhN^Aδ-LTMR^ inhibitory populations. Each IhN subtype is named by its dominant sensory input. **b**, Representative skeleton reconstructions of IhN^Aδ-HTMR^, islet^C-HTMR/Heat^, islet^C-LTMR^, and islet^Aδ-LTMR^ neurons. Axons are shown in black. **c**, Summary of dendro-dendritic synapses from inhibitory to excitatory subtypes. Dot size indicates mean synapse count for each subtype pair; color indicates mean synapse density per unit overlap (capped at the 95th percentile). **d**, IhN inputs onto the ExN and PN subpopulations (dendro-dendritic synapses only). Islet-PSI-IhN^C-HTMR/Heat^ and Islet-PSI-IhN^C-LTMR^ uniquely form dendro-dendritic synapse. **e,** Same as c, but for axo-dendritic synapses. **f,** Same as d, but for axo-dendritic synapses. **g**, Schematic illustrating heterotypic feedforward inhibition in the dorsal horn. Both C-LTMRs and Aδ-HTMR evoke feedforward inhibition to C-HTMR/Heat modules.

**Figure S9.**
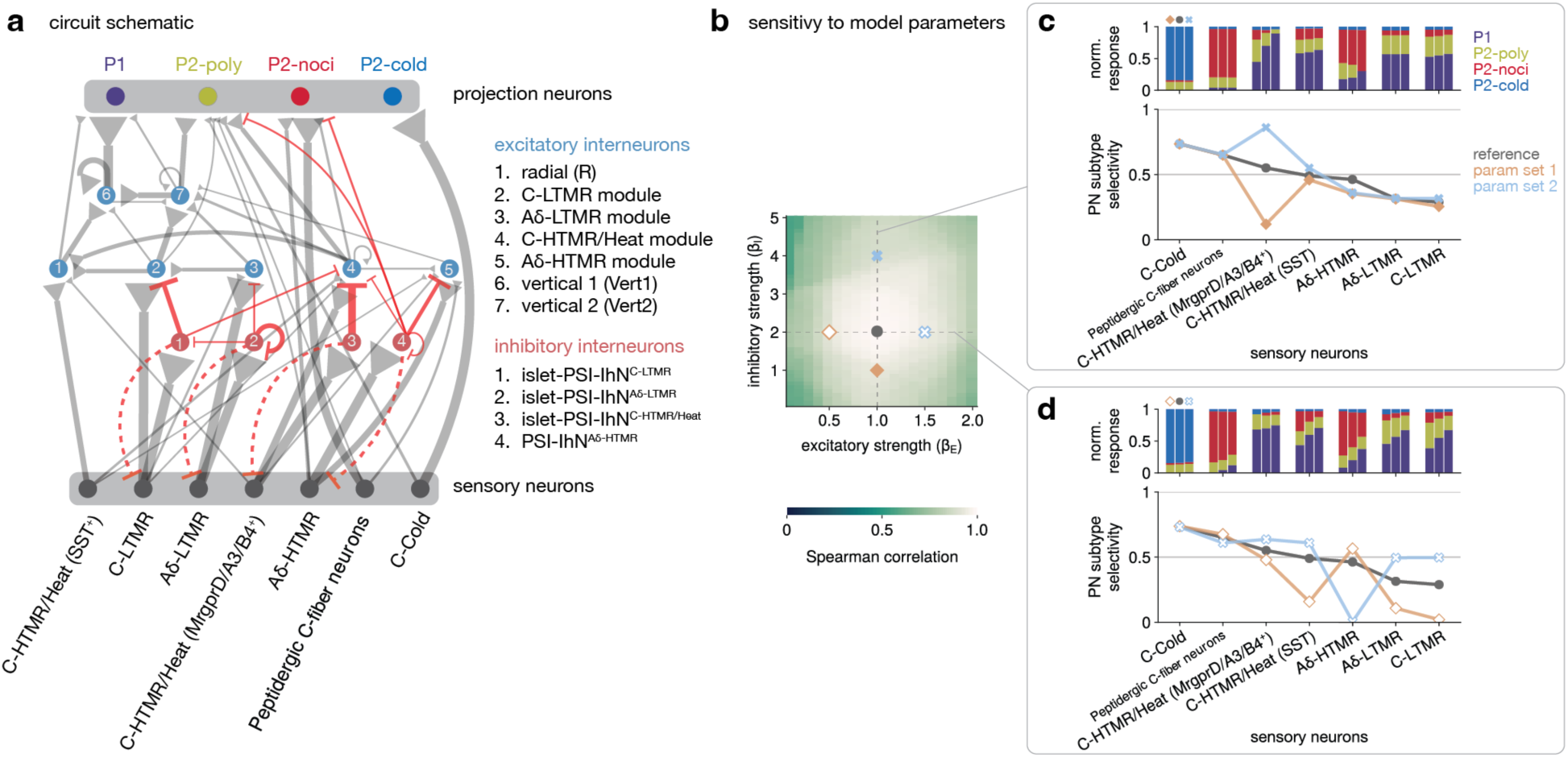
Wiring diagram of the model and sensitivity to model parameters. **a**, Schematic of dorsal horn circuit derived from reconstructed sensory afferents and dorsal horn neurons. Each node represents a distinct neuronal population: sensory afferents (gray), excitatory interneurons (ExNs, blue), inhibitory interneurons (IhNs, red), and projection neurons (PNs, colored). Edge thickness between nodes is proportional to the corresponding weight in the connectivity matrices of Figure 5a, with weights capped at 0.5 and edges with weights < 0.065 omitted from the schematic for clarity; thus, the thickest edges correspond to a weight of 0.5 and the thinnest to 0.065. These weights reflect the relative contribution of each presynaptic population to the synaptic input of a typical postsynaptic neuron (of a given type) in the model (see Methods) and should be interpreted differently from the wiring diagram in Figure 4, where arrow thickness represents average synapse counts. Axoaxonic synapses mediating presynaptic inhibition are depicted as dashed lines, with thickness not scaled to synapse count, and are excluded from the model for simplicity. Note the strength of feedforward inhibitory motifs between sensory afferents, islet neurons and their corresponding ExN modules. Also note the strong direct pathways between C-Cold and peptidergic C-fiber neurons and their targets in the output (PN) layer. **b**, Sensitivity of the model to parameters, specifically the strengths of excitation (*β_E_*) and inhibition (*β_I_*). *β_E_* scales the excitatory weights originating from ExNs, and *β_I_* scales the inhibitory weights (see Methods). The heatmap shows the Spearman correlation between sensory-evoked normalized PN responses at the reference parameter set (*β_E_* = 1, *β_I_* = 2; gray circle) and those obtained for different (*β_E_*, *β_I_*) pairs. **c**, Analogous to Figure 5c, but with three values of *β_I_*: filled diamond, circles (reference), and filled cross, with *β_E_* fixed at the reference value. Normalized PN responses for each parameter set are shown as stacked bars (the center stacked bar corresponds to the reference parameter set). The corresponding selectivity metric is shown in the lower panel. Note that the identity of the preferred PN subtype is preserved across a wide range of parameter choices, despite changes in the degree of selectivity. The sensitivity of PN responses to *β_I_* differs across sensory stimulation conditions (see Methods). **d**, Same as in **c**, except three values of *β_E_*, with *β_I_* fixed at the reference value.

## Methods

### Mice

All mice used in this study were maintained on a mixed background. Mice were handled and housed in accordance with Harvard Medical School and IACUC guidelines. Mice were kept in a temperature- and humidity-controlled room with a 12-hour light/dark cycle, with food and water available *ad libitum*. A postnatal day 100 (P100) female *Th^T2A-CreER^; ROSA26^LSL-Matrix-dAPEX2^* mouse was used to generate the EM dataset. C-LTMRs were labeled in this EM volume by tamoxifen administration (1 mg/day) at P21–P23 (Abraira et al., 2017). Grpr^+^ vertical cells were labeled by crossing *Grpr^CreER^* mice with Ai14 mice and administering tamoxifen (0.5 mg) at P42 (Polgar et al., 2023). NPFF^+^ vertical cells were labeled by injecting AAV2/8-hSyn-DIO-mCherry into the spinal dorsal horn of *NPFF^Cre^* mice at P21 (Quillet et al., 2023). Gpr83^+^ lamina I projection neurons were labeled by crossing *Gpr83^CreER^; Phox2a^FlpO^* mice with Ai65 mice and administering tamoxifen (1.25 mg) at P42 (Choi et al., 2020). Tacr1^+^ projection neurons were labeled by crossing *Tacr1^CreER^; Phox2a^FlpO^* mice with Ai65 mice and administering tamoxifen (0.5 mg) at P42 (Choi et al., 2020). Generation and characterization of the *Phox2a^FlpO^* will be described elsewhere.

### Tamoxifen administration

Tamoxifen (MilliporeSigma) was dissolved in ethanol (20 mg/mL), mixed 1:1 with sunflower seed oil (MilliporeSigma), and vortexed. Ethanol was then removed under vacuum, and the resulting solution was administered to animals by intraperitoneal or oral injection.

### Immunochemistry

Mice were anesthetized with a high dose of inhaled isoflurane and transcardially perfused with 15 mL of ice-cold 1X PBS and 25 mL of 4% PFA in 1X PBS. Spinal columns were dissected out and post-fixed in 4% PFA rotating at 4°C overnight, then washed with 1X PBS 3 times for 10 minutes. Spinal cords were microdissected and placed in 30% sucrose at 4°C for 1-2 nights or until the tissue sank. Tissues were then placed in cryomolds in Epredia™ Neg-50™ Frozen Section Medium (Fisher, 22-046-511) and frozen over dry ice. Sagittal sections of lumbar spinal cord were cryosectioned at 250 μm and collected into 24-well plates containing PBS. All steps were performed with gentle agitation. Sections were washed in 0.3% TritonX-100 in PBS 5 times for 30 minutes, then transferred to blocking solution (5% normal donkey serum, 0.3% TritonX-100 in PBS) for 2 hours at room temperature. Slices were then incubated in blocking solution with primary antibodies (goat anti-mCherry, Origene, AB0040-200, 1:500; rabbit anti-CGRP, Immunostar, 24112, 1:500) for 72 hours, washed as described above and incubated in blocking solution with secondary antibodies (donkey anti-goat Alexa Fluor™ 546, Thermo Fisher, A11056, 1:1000; donkey anti-rabbit Alexa Fluor™ 488, Thermo Fisher, A32790, 1:1000; IB4 Alexa Fluor 647 conjugated, Thermo Fisher, I32450, 1:500) for 48 hours. Slices were washed again and then dehydrated with serial incubation in 50%, 75%, and 100% methanol in PBS at room temperature for 10 minutes each and kept in 100% methanol overnight for full dehydration. Before imaging, slices were cleared in BABB solution (1 part Benzyl Alcohol, 2 parts Benzyl Benzoate) for 2 hours. Sections were imaged using Zeiss LSM-900 confocal microscope at 10X magnification for general morphology and 63X magnification for dendritic spine quantification.

### EM sample preparation

EM sample preparation was performed as described previously (Xiang et al., 2026a). Briefly, the animal was transcardially perfused with aldehyde fixative. Lumbar spinal cord segments L3–L4 were dissected and sectioned sagittally at 200 µm. After DAB staining, sections were post-stained using the rOTO protocol, dehydrated, and embedded in resin.

### EM volume generation

EM volume generation was performed as described in a parallel study (Xiang et al., 2026a). Serial-section TEM data were acquired using the GridTape pipeline (Phelps et al., 2021). Following imaging, EM images were segmented using convolutional neural networks (CNNs). Synapses were automatically predicted using CNNs that detect synaptic clefts and assign pre- and postsynaptic objects (Buhmann et al., 2021; Lee et al., 2017).

### Targeted neuron proofreading

We focused reconstruction and proofreading primarily on spinal cord excitatory interneurons in laminae I-II. A region of interest (ROI) centered within the EM volume was selected, spanning 230 µm × 110 µm × 31 µm along the rostral–caudal, dorsal–ventral, and medial–lateral axes, respectively. Laminae I-II were defined dorsoventrally from the white matter border to the ventral boundary of the APEX-labeled C-LTMR terminal field.

Excitatory and inhibitory neurons were distinguished based on ultrastructural criteria. Excitatory neurons were identified by the presence of asymmetric synapses with round, clear vesicles, whereas inhibitory neurons were identified by the presence of symmetric synapses with vesicles of variable size and shape (**Fig. S1j,k**). In addition, many inhibitory neurons in laminae I-II formed axoaxonic synapses onto sensory terminals and/or dendrodendritic synapses onto other neurons, consistent with known features of GABAergic dorsal horn interneurons. Three neurons were excluded from our analysis because they formed both symmetric and asymmetric synapses. Using these criteria, we identified 122 excitatory interneurons (including 22 vertical cells), 33 inhibitory interneurons, and 2 projection neurons within the densely reconstructed ROI.

We then densely reconstructed all 122 excitatory neurons. Proofreading was performed using Neuroglancer and CAVE (Dorkenwald et al., 2025). Dendritic segmentation quality was high; for each neuron, we verified all major branch points to ensure accurate reconstruction and, when necessary, split falsely merged dendritic segments. For axons, we proofread each process to its terminal tip or to the point at which it exited the imaged volume.

For characterizing feedforward excitatory circuits onto projection neurons, we additionally reconstructed projection neurons and vertical cells outside the ROI, since vertical cells provide presynaptic input to projection neurons. Therefore, in total, we reconstructed 77 vertical cells and 46 projection neurons spanning the full EM volume. For projection neurons, we proofread dendritic arbors only. The reconstructed ExNs had a total of 0.964 m of axon length, with individual neurons having an average of 5.6 mm of axon length.

### ExN feature construction and clustering

Historically, spinal cord interneurons have been classified primarily by dendritic morphology (e.g., radial vs. vertical cells) and their laminar position. Here, we developed a clustering workflow that instead groups neurons using a combined feature space integrating dendritic morphological metrics and primary sensory neuron input composition, allowing us to identify neuron classes that reflect both structure and synaptic connectivity.

For each neuron, we built a unified feature vector by concatenating (i) dendrite morphology features computed from SWC skeletons and (ii) sensory-input features computed from synapse counts.

1. Dendrite morphology features (SWC-based).

SWC files were parsed to extract nodes and parent–child edges. We computed dendrite-only summary statistics (axon-related metrics were computed but excluded from clustering features). The morphology feature set included:

- Size/complexity: total dendrite cable length (dend_len_um), dendritic tip number (tips_dend), dendritic branch point number (branch_dend), length-normalized measures (branch_per_100um, tips_per_100um), and mean dendritic segment length (mean_seglen_dend).
- Reach/extent: maximum dendritic path length from soma (max_path_dend_um), 3D span (x_span_um, y_span_um, z_span_um), and bounding-box volume (bbox_vol_um3).
- Geometry/orientation: dendritic directionality (directionality_dend, based on the mean unit vector of dendritic edges).
- Soma position: soma depth along the y-axis (soma_y_um).

2. Sensory-input features (synapse-count-based).

We loaded a per-neuron sensory synapse count table and used the following set of sensory categories: Aδ-HTMR, C-Cold, C-HTMR/Heat (Sst+), C-HTMR/Heat (MrgprD+, MrgprB4+, MrgprA3+), Peptidergic C-fiber neurons, Aβ-LTMRs, Aδ-LTMRs, C-LTMRs, low-confidence sensory predictions, and non-sensory inputs. From these counts, we derived sensory features:

- Fractional inputs: each category divided by the neuron’s total synapse count.
- Log-transformed counts: log1p(count) per category.
- CNS fraction: non-sensory synapses /total synapses to capture non-sensory (CNS) input proportion.

For the 177 neurons, morphology and sensory tables were aligned by neuron ID and concatenated into a single 177 × 33 matrix (neurons × features).

We standardized features using column-wise z-normalization, embedded neurons into 2D with UMAP (Euclidean metric; n_neighbors=5, min_dist=0.21), and then clustered neurons in the 2D UMAP space using HDBSCAN (McInnes, 2018).

After unsupervised clustering, a human expert (W. X) reviewed each cluster and finalized class labels based on dendritic morphology and primary sensory input profiles. Only 3 of 177 neurons required label changes during expert curation, indicating strong agreement between the automated clustering and experts.

### PN clustering

To identify lamina I projection neurons (PNs), we leveraged previous reports that PNs have larger somata and more elaborate dendritic arbors than lamina I interneurons. For each lamina I neuron, we cropped the soma mesh within a 15 μm radius of a manually defined soma center and repaired the cropped mesh to generate a watertight surface for reliable volume estimation. We focused on 456 lamina I neurons that received at least one synapse from reconstructed excitatory interneurons (ExNs). We computed their soma volumes and observed a bimodal distribution consistent with populations of small interneurons and large PNs (**Figure S3a**). Based on this distribution, 61 neurons with soma volumes >1,190 μm³ were classified as PN candidates. We then manually inspected all 61 candidates and excluded 12 vertical cells, one islet cell, and three neurons with less elaborate dendritic arbors. We additionally included one projection neuron with a partially reconstructed soma located at the edge of the EM volume. This yielded a final dataset of 46 putative projection neurons for subsequent analyses. These putative PNs comprise approximately 10% of lamina I neurons, consistent with previous estimates that projection neurons account for 5–10% of the lamina I neuronal population (Al Ghamdi et al., 2009; Bice and Beal, 1997a, b).

We next stratified PNs using a combination of morphology and sensory input profiles. We observed that 6 PNs had ventrally directed dendrites extending into the LTMR recipient zone, consistent with reported properties of Gpr83*^+^* PNs and with our sparsely labeled reconstructions. We therefore designated these 6 neurons as P1 (putative Gpr83*^+^* PNs) and excluded them from subsequent input-based clustering. The remaining 40 PNs had dendritic arbors restricted above lamina IIid, consistent with Tacr1+ PN morphologies; we refer to this set as P2 (putative Tacr1*^+^*). We used SPINE to quantify sensory inputs to both P1 and P2 neurons and found that P1 neurons receive very sparse monosynaptic sensory input (∼3% of total inputs), in striking contrast to P2 neurons, which receive substantially more direct sensory input. We then clustered P2 neurons based on their sensory synaptic input profiles (**Fig. S3**). Because many PNs were truncated at the volume boundaries, we performed UMAP embedding and HDBSCAN clustering using only synaptic input features (rather than morphology plus input features, as in the ExN analysis). To emphasize selectivity for nociceptive drive, we combined Aδ-HTMR and peptidergic C-fibers inputs into a single nociceptor category prior to embedding and clustering. All ExNs and PNs cell type label after clustering are available as a CAVE table named ‘spinalcord_neuron_clusters_2’.

### Sensory input analysis

For all spinal cord neurons, we first queried their input synapses and presynaptic neurons from CAVE (synapse table named ‘synapses_version3’). We then embedded the presynaptic neuron cutout using the SPINE model as described in the accompanying manuscript (Xiang et al., 2026a). A presynaptic neuron cutout was classified as a CNS neuron, a specific sensory subtype, or low-confidence sensory neuron if sensory subtype scores were ≤ 0.6.

### Path overlap calculation

To quantify spatial overlap between a presynaptic neuron process (axon or dendrite) and a postsynaptic neuron process (axon or dendrite), we measured the geodesic extent along the presynaptic skeleton that lies within a fixed Euclidean 3D distance of the postsynaptic skeleton. Neurons were skeletonized using the pcg_skel pipeline (pcg_skeleton). Neuronal compartments (axon vs dendrite) were assigned using pcg_meshwork, which infers an is_axon annotation from synapse directionality (pre-vs post-synapses). This axon annotation is defined on the mesh but was mapped onto skeleton vertices via the skeleton–mesh correspondence, enabling selection of axon- or dendrite-labeled skeleton vertex subsets.

For each pre–post pair and chosen compartment combination, we extracted 3D coordinates of the corresponding presynaptic and postsynaptic skeleton vertices. Prior to overlap analysis, each skeleton was resampled along every graph edge so that consecutive vertices along an edge were placed at approximately uniform spacing: edges were subdivided with a target internode distance of 100 nm (short edges retain only their endpoints and therefore can be shorter than 100 nm). We then built a scipy.spatial.cKDTree on the postsynaptic vertex coordinates and queried each presynaptic vertex for neighbors within 5 µm, yielding a set of close vertex pairs (v_pre_, v_post_). Close pairs were grouped into contiguous overlap regions using skeleton connectivity in both neurons: we constructed adjacency lists from each skeleton’s edge list and clustered pairs such that presynaptic vertices were connected within one graph hop on the presynaptic skeleton and postsynaptic vertices were connected within one graph hop on the postsynaptic skeleton. For each overlap region, we computed the geodesic diameter (maximum shortest-path distance) on the induced subgraph of presynaptic skeleton vertices in that region, using Dijkstra shortest paths with edge weights equal to the Euclidean lengths of skeleton edges. The resulting geodesic diameter (reported in µm) was taken as the path overlap length along the presynaptic process.

### Probabilistic network traversal analysis

To organize excitatory interneurons (ExNs) and projection neurons (PNs) according to their position within sensory pathways, we applied a probabilistic network traversal model adapted from (Schlegel et al., 2021) and implemented using navis.models.TraversalModel. We constructed a directed, weighted connectivity matrix containing synaptic connections from sensory afferents and ExNs onto ExNs and PNs. Aβ low-threshold mechanoreceptor afferents were excluded from this analysis. All connections containing at least one synapse were retained, with edge weights corresponding to the number of synapses between each presynaptic and postsynaptic neuron. The resulting connected network contained 135 sensory afferents, 177 ExNs and 46 PNs, comprising 6,506 connections and 24,947 synapses. One C-Cold afferent that formed no connections within the network was excluded.

For each connection from presynaptic neuron *i* to postsynaptic neuron *j*, the normalized input fraction was calculated as

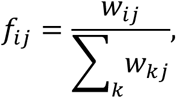

where *w_i_*_j_is the number of synapses from neuron *i* onto neuron *j*, and the denominator is the total number of synapses received by neuron *j* from all sensory afferents and ExNs included in the network. Thus, input fractions represent the relative contributions of reconstructed sensory and excitatory inputs and do not include inhibitory or untraced inputs.

The 135 connected sensory afferents were used as the seed population and assigned to layer 1. At each subsequent step, connections from previously traversed neurons to untraversed postsynaptic neurons were independently evaluated using a linearly scaled traversal probability,

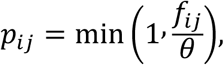

where *θ*is the traversal threshold. We used *θ* = 0.5, such that a connection contributing at least 50% of the reconstructed sensory and excitatory input to its target was always traversed, whereas weaker connections were traversed with probabilities proportional to their input fractions. The threshold was used only to determine traversal probability and was not used to exclude connections. A postsynaptic neuron was traversed when at least one eligible incoming connection was successfully traversed. Connections onto neurons not reached during a given step were evaluated again at subsequent steps, allowing weakly connected neurons to be reached later than neurons receiving strong or direct sensory input.

The traversal was repeated for 10,000 Monte Carlo iterations using a fixed NumPy random seed, with a maximum of 50 steps per iteration. All neurons were reached in every iteration, and the maximum observed first-encounter step was 11, indicating that the step limit did not truncate the analysis. The traversal layer of each neuron was defined as the mean step at which it was first reached across iterations. This measure therefore represents a probabilistic first-encounter distance from the sensory periphery rather than the shortest-path distance through the network.

Cell-type layer values were calculated by averaging the resulting neuron-level traversal layers within each ExN or PN subtype.

To assess robustness to the traversal threshold, the analysis was repeated using *θ* = 0.05, 0.1, 0.2, 0.3, 0.5, 0.75 and 1.0, with 2,000 iterations per threshold. Changes in the ordering of cell types were evaluated using Spearman rank correlations between their mean traversal layers across threshold values.

### Construction of the wiring diagram

To generate the wiring diagram linking sensory neuron, ExN, PN subtypes, we quantified the mean synapse number and mean synapse density for each presynaptic–postsynaptic cell-type pair from reconstructed neurons. Connections with a mean synapse count <0.25 synapses and a mean synapse density <0.036 synapses per 10 μm of axon–dendrite path overlap were excluded. For sensory afferent inputs, we additionally incorporated the comprehensive SPINE sensory input analysis, which classifies all sensory synapses onto reconstructed neurons. Sensory neuron– ExN/PN connections contributing, on average, less than 3.5% of the total synaptic input to the postsynaptic population were excluded. Three sensory connections exceeded the 3.5% input threshold but fell below the synapse count and/or synapse density thresholds; these connections were retained in the wiring diagram and displayed as dashed lines.

In the final wiring diagram, line thickness represents the mean synapse number between cell-type pairs: 0.5 pt (0.25–0.5 synapses), 1 pt (0.5–1 synapse), 2 pt (1–2 synapses), 4 pt (2–4 synapses), and 8 pt (>4 synapses). Connections with <0.25 mean synapses are shown as 0.3 pt dashed lines. Line color represents mean synapse density: dark blue (>0.14 synapses per 10 μm path overlap), medium blue (0.07–0.14), light blue (0.036–0.07), and gray dashed (<0.036). Aβ-LTMRs are not shown because they contribute, on average, less than 3.5% of the total synaptic input to the reconstructed spinal cord neuron types included in this diagram.

### Connectome-constrained model of the dorsal horn microcircuit

The connectome of the dorsal horn circuitry consists of two components: a feedforward component, representing the connectivity between sensory afferents and dorsal horn neurons, and a recurrent component, representing the local connections within the dorsal horn. Neurons in the dorsal horn include excitatory (ExNs) and inhibitory interneurons (IhNs) as well as projection neurons (PNs). Our goal was to construct a model that would allow us to assess the effect of sensory subtype activations on downstream dorsal horn neurons, particularly on PNs. Due to the low recall rate for automated axoaxonic synapse prediction, presynaptic inhibition mediated by axoaxonic IhN–afferent connections were not incorporated into the model.

We consider a firing rate-based model of the form

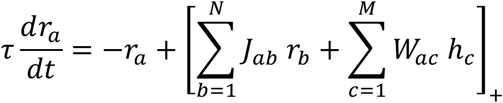

where *r_a_* denotes the firing rate of dorsal horn population *a*, with *a*, *b* = 1, …, *N*. Similarly, ℎ*_c_* denotes the firing rate of sensory population *c*, with *c* = 1, …, *M*. In this model, each unit, *r_a_*, represents the population-averaged activity of one dorsal horn neuron subtype, and each ℎ*_c_* the corresponding sensory neuron subtype. The recurrent and feedforward contributions are given by ***Jr*** and ***Wh***, respectively. The operation [⋅]_+_ denotes rectified linear thresholding, which ensures nonnegative firing rates. The network time constant *τ* is set to 1, so that time is measured in units of *τ*.

In the model, dorsal horn neurons consist of excitatory interneurons (radial, C-LTMR module, Aδ-LTMR module, C-HTMR/Heat module, Aδ-HTMR module, Vert1, and Vert2), inhibitory interneurons (Islet-PSI-IhN^C-LTMR^, Islet-PSI-IhN^C-HTMR/Heat^, Islet-PSI-IhN^Aδ-LTMR^, and PSI-IhN^Aδ-HTMR^), and projection neurons (P1, P2-poly, P2-noci, and P2-cold). Sensory neurons comprise SST^+^ C-M/H, C-LTMR, Aδ-LTMR, MrgD/A3/B4^+^ C-M/H, Aδ-HTMR, peptidergic C-fibers, and C-cold. Throughout this section, we use the term ‘subtype’ to refer to each of these individual neuron classes.

The model is constrained by the connectome in that the connectivity matrices ***J*** and ***W*** are derived from empirical synapse counts between fully reconstructed neurons. Because the model abstracts each subtype as a single representative neuron, the synaptic connectivity between any two subtypes must be reduced to a scalar. To do so, we began with the raw feedforward and recurrent synaptic connectivity matrices, denoted by 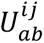 and 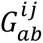, respectively. In this notation, *a*, *b* index subtypes, whereas *i*, *j* index individual cells within a given subtype.

We first accounted for the variability in the total number of synaptic inputs across reconstructed neurons. To do so, we calculated the total in-degree for each dorsal horn neuron from 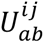 and 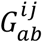. The in-degree, defined as the sum of synapse counts converging onto neuron *i* in subtype *a*, is

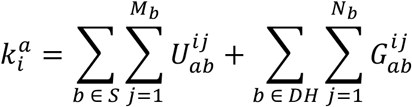

where *S* and *DH* denote the sets of sensory and dorsal horn neuron subtypes listed above, respectively. *M_b_* is the number of cells within sensory subtype *b*, and likewise *N_b_* for dorsal horn subtype *b*. We then defined the in-degree-normalized connectivity matrices, or matrices of ‘fractional synapse counts’, by 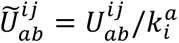, and 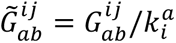. In this normalization, the strength of a given synapse in driving a postsynaptic response depends on the total synapses the postsynaptic neuron receives, which accounts for sampling biases in the reconstruction process and the natural variability in total synaptic input across neurons.

Next, we constructed ***J*** and ***W*** by

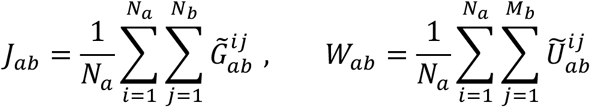

In other words, for a pair of subtypes, *b* (presynaptic) and *a* (postsynaptic), we computed the total fractional synapse count and then normalized by the size of the postsynaptic population, *N_a_*. This normalization yielded an effective synaptic strength per postsynaptic neuron in subtype *a*. In this formulation, a larger presynaptic population produces a stronger postsynaptic drive than a smaller presynaptic population, independent of the size of the postsynaptic population.

Lastly, the signs of the columns of ***J*** are constrained, such that columns corresponding to inhibitory subtypes are multiplied by -1. Further, excitatory and inhibitory columns in ***J*** are scaled by *β_E_* and *β_I_*, respectively, to capture potential differences in the strength of excitatory and inhibitory synapses. Columns in ***W*** remain non-negative. In **Figure 5**, we set *β_E_* = 1 and *β_I_* = 2; **Figure 5a** shows the resulting heatmaps of ***J*** and **W**.

We further explored the effects of varying *β_E_* and *β_I_* (**Figure S8b-d)**. We found that the identity of the preferred PN subtype was preserved across a wide range of parameter choices. We also found that increasing the strength of inhibition suppressed the response of P2-poly PNs to stimulation of C-HTMR/heat (MrgprD/A3/B4^+^) or Aδ-HTMR afferents, but left its response to Aδ-LTMR, Peptidergic C-fibers, and C-Cold stimulation largely unchanged. More generally, the sensitivity of PN responses to *β_E_* and *β_I_* differs across sensory stimulation conditions and reflects the relative contributions of excitatory and inhibitory pathways onto the respective PN subtypes. The sensory input ***h***(*t*) takes the form of a step function, ***h***(*t*) = ***h*_0_** during stimulation and ***h***(*t*) = **0** otherwise, where ***h*_0_** represents the activation pattern across sensory afferent subtypes. In the stimulation experiments, we considered a binary ***h*_0_**, where all entries are set to zero except one, corresponding to the sensory subtype of interest. This isolates the effect of a specific sensory subtype on downstream neurons, particularly on the four PNs (**Figure 5b-c**). We also consider non-binary patterns in ***h*_0_** (see below). The model was numerically integrated using the Euler method with simulation time-step of 0.01, and the stimulation duration was sufficient to ensure the firing rates reached a steady state.

Next, we considered a case where ***h*_0_** is not binary and is instead optimized to produce the largest steady-state response in a given PN. This reveals the sensory neuron activity pattern that is ideally suited to drive a particular PN. In this optimization, we impose the constraints ‖***h*_0_**‖_2_ = 1 and 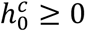 for all sensory neuron subtypes *c*, and define the objective 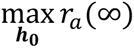, where *a* indexes the PN of interest. The optimization was performed using the Nelder-Mead method (100 random initializations), implemented with the Optim.jl package in Julia.

The selectivity index in Figure 5c is calculated as

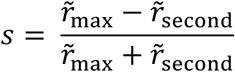

where r̃_max_ and r̃_second_ denote the largest and second-largest values of r̃*_a_* across the four PN subtypes, respectively, with r̃*_a_* denoting the normalized steady-state response of the *a*^th^ PN subtype, namely r̃*_a_* = *r_a_*(∞) / ∑*_b_ r_b_*(∞). The metric *s* ranges from 0 to 1: *s* = 1 when only the most-activated PN subtype has a nonzero response, and *s* = 0 when the two most-activated PNs are equally activated.

From simulation results, we highlight three qualitative agreements between individual sensory neuron stimulations in the model and optogenetic sensory perturbation experiments in Zhang et al., 2024 and Choi et al., 2020. In these experiments, PNs are identified by their gene expression patterns: Gpr83^+^ corresponds to P1, while Tacr1^+^ labels both P2-poly and P2-noci PNs.

First, model stimulation of C-LTMR (**Figure 5b**) agrees with Figure 4c of Zhang et al., 2024, showing comparable levels of activity in P1 and P2-poly. Second, Aδ-HTMR stimulations in the model (**Figure 5b**) are consistent with the optogenetic stimulation experiments in Figure 3k of Choi et al., 2020, in which P2-noci, but not P1, exhibits a pronounced response. Third, stimulating C-HTMR/Heat (MrgprD/A3/B4^+^) in the model (**Figure 5b**) produces comparable responses in P1 and P2-poly, in agreement with Extended Data Figure 7f of Choi et al., 2020.

## Acknowledgements

The authors thank T. Docter, A. Shuster, S. Deng, S. Markman, LF Abbott, M. Beiran, K. Ng and members of the Ginty and Lee laboratories for discussions and comments on the manuscript; A.J. Todd for sharing the *Grpr^CreER^* and *NPFF^Cre^* mouse lines; J. Joroff, E. Yacharn, E. Xue, K. Ohgami, S. Simhadri for tracing neurons; the CAVE team for managing and maintaining the CAVE servers that host the dataset; S. Gerhard for advice on analyzing neuron skeleton; F. Collman, C.S.-Mizell for advice analyzing neuron mesh data; J. Phelps, A.T. Kuan, D.G.C. Hildebrand, B. Graham, K. Kris for advice on EM imaging and alignment; R. Schalek for performing micro-CT scans on the EM samples; HMS IT for assisting with data storage; L. Qi for assistance with animal surgery and breeding. The authors gratefully acknowledge BossDB.org for providing data storage and access services that support the sharing of the datasets associated with this publication (R24MH114785, Hider et. al 2022 – https://doi.org/10.3389/fninf.2022.828787). We thank Caiying Guo and the Janelia Gene Targeting and Transgenic Facility for generating the *Phox2a^FlpO^* mouse line. This work was supported by Stuart H.Q. and Victoria Quan fellowships (W.X., J.L. and Q.Z.), NIH grant R35 5R35NS132196 (D.D.G.), the K. Lisa Yang Brain Body Center at Harvard Medical School (D.D.G.), the Edward R. and Anne G. Lefler Center for Neurodegenerative Disorders (D.D.G.), and an R&D Impact Foundation Neurobiology Spark Grant (W.C.A.L.). D.D.G. is an investigator of the Howard Hughes Medical Institute.

## Author contributions

W.X., D.D.G. and W.-C.A.L. conceptualized the project and designed experiments. W.X. generated the EM dataset. K.D. and W.X. developed and applied the SPINE model. T.N., T.M. and N.K. developed and applied the segmentation pipeline. J.A.B. developed and applied the synapse prediction pipeline. A.H. set up CAVE for reconstruction. W.X., S.C., Q.Z., B.M.L., A.S., M.F.M., J.N., R.B., L.S.C., R.W., A.D., A.N. reconstructed the neurons. W.X., K.D., R.K., A.H., S.W., D.Z., P.L.T analyzed the connectivity data. R.K. performed circuit modeling. J.L. and W.X. performed the sparse labeling experiments. S.C. helped characterize the *Phox2a^Flpo^*line. W.X., D.D.G. and W.-C.A.L. wrote the manuscript with input from other authors.

## Data availability

The volumetric EM datasets generated in this study, including the dSC1 dataset and dSC_APEX datasets, segmentation, and cell-type annotations are open source and publicly available through: https://github.com/htem/spinal_cord_connectome.

## Code availability

Code for the development and application of SPINE is available at: https://github.com/htem/dorsalhorn-ml. Code for analysis and figure generation is available at: https://github.com/htem/spinal_cord_connectome.

